# Improving the Resolution and Informativity of the Majority-Rule Consensus Tree Using Fine-Grained Dissimilarity Measures

**DOI:** 10.64898/2026.03.16.712085

**Authors:** Yuki Takazawa, Atsushi Takeda, Momoko Hayamizu, Olivier Gascuel

## Abstract

Phylogenetic analyses often require the summarization of multiple trees, e.g., in Bayesian analyses to obtain the centroid of the posterior distribution of trees, or to determine the consensus of a set of bootstrap trees. The majority-rule consensus tree is the most commonly used. It is easy to compute and minimizes the sum of Robinson-Foulds (RF) distances to the input trees. In mathematical terms, the majority-rule consensus tree is the median of the input trees with respect to the RF distance. However, due to the coarse nature of RF distance, which only considers whether two branches induce exactly the same bipartition of the taxa or not, highly unresolved trees can be produced when the phylogenetic signal is low. To overcome this limitation, we propose using median trees with respect to finer-grained dissimilarity measures between trees. These measures include a quartet distance between tree topologies, and transfer distances, which quantify the similarity between bipartitions, in contrast to the presence/absence view of RF. We describe fast heuristic consensus algorithms for transfer-based tree dissimilarities, capable of efficiently processing trees with thousands of taxa. Through evaluations on simulated datasets in both Bayesian and bootstrapping maximum-likelihood frameworks, our results show that our methods improve consensus tree resolution in scenarios with low to moderate phylogenetic signal, while providing better or comparable dissimilarities to the true phylogeny. Applying our methods to mammals, Ebola virus, and a large HIV dataset of over nine thousand taxa confirms the improvement with real data. These results demonstrate the usefulness of our new consensus tree methods for analyzing the large datasets that are available today. Our software, PhyloCRISP, is available from https://github.com/yukiregista/PhyloCRISP.

## Introduction

Phylogenetic analysis is a fundamental tool in computational biology for understanding the evolutionary relationships among taxa. In many scenarios, such as Bayesian analysis or bootstrap analysis, the result is not a single phylogenetic tree but a set of trees.

There are two common ways to use these sets of phylogenetic trees. One is to use this set to assign support values to each branch in some tree estimate, indicating how robust or credible it is. Researchers typically “trim” or simply ignore those branches with low support values when drawing conclusions. The most standard support value is the frequency of occurrence of each branch (more precisely, the bipartition or split of the taxa induced by the given branch) in the input trees. This corresponds to the Felsenstein bootstrap proportion (FBP; Felsenstein 1985). The other approach is to use a consensus method that provides a single tree to capture the central tendency of the tree distribution. This approach seems natural when dealing with Bayesian posterior tree draws, since the posterior distribution itself is the object of estimation and there is no reference tree estimate. The use of a consensus tree to summarize the bootstrap tree distribution is the approach that was taken in Felsenstein’s original CONSENSE program (Felsenstein 1989), over the alternative of considering the reference tree with branch supports.

Many approaches have been developed since Adams consensus tree (Adams 1972). Bryant (2003) provides a summary and a classification of these methods. To construct a consensus of unweighted, unrooted trees with the same set of taxa, most of the popular methods use either bipartition-based or quartet-based approaches. Bipartition-based methods find a tree-compatible set of bipartitions from the input trees. Such methods include the strict consensus tree, the majority-rule consensus tree, the loose consensus tree (Bremer 1990), the extended majority-rule consensus tree (also named the greedy consensus tree), and the Nelson-Page consensus tree (Nelson 1979; Page 1990). The strict and majority-rule consensus trees tend to be poorly resolved, while the extended version is typically highly resolved, although it is not guaranteed to be fully resolved (i.e., binary, where every non-leaf node has degree 3). Quartet-based methods consider quartet topologies, where a quartet topology (or quartet for brevity) refers to a tree topology restricted to four taxa. Each quartet can take one of the four possible forms: three resolved topologies with one internal branch and one unresolved “star” topology. Quartet-based methods include the Q* consensus tree (Berry and Gascuel 2000) and the weighted quartet distance consensus (WQDC; Yourdkhani and Rhodes 2020). ASTRAL (Mirarab et al. 2014; Zhang et al. 2025) is a quartet-based method that was developed for the problem of species tree reconstruction from a set of gene trees. ASTRAL and the programs of the ASTRAL family always output a fully resolved tree, while Q* trees are generally poorly resolved. Although ASTRAL is not often used for consensus, as long as the limitation that it produces fully resolved trees is clear, it can be used as a median tree, based on quartet tree topologies (Arasti and Mirarab, 2024).

In Bayesian phylogenetics, in addition to traditional consensus trees, two methods are frequently used to generate a single tree to summarize the set of posterior trees drawn by Markov chain Monte Carlo (MCMC) methods. The first one is the maximum *a posteriori* (MAP) tree, which is the highest peak of the posterior distribution. The MAP tree practically refers to one of the posterior trees drawn by MCMC that has the maximum posterior value. The second one is the maximum clade credibility (MCC) tree, which is the default consensus tree output produced by TreeAnnotator in both BEAST1 (Suchard et al. 2018) and BEAST2 (Bouckaert et al. 2019) and is defined as one of the input trees with the largest sum of branch posterior probabilities. Although these two methods do not combine information from multiple input trees to create a new tree topology, they share the goal of summarizing the input tree distribution. For simplicity, we also refer to them as consensus trees in this paper. Recently, proper consensus alternatives to those two methods have been proposed. These include HIPSTR (Baele et al. 2025) and CCD-MAP (Berling et al. 2025) trees. Note that all these consensus trees (MAP, MCC, HIPSTR, and CCD-MAP) are fully resolved.

Of all the consensus trees discussed above, the majority-rule consensus tree is probably the most commonly used method for summarizing a set of trees. The majority-rule consensus tree is defined by the bipartitions that appear in more than half of the input trees and can be seen as an intermediate balanced approach between the highly conservative strict consensus tree and the more inclusive extended majority-rule tree. The strict consensus tree retains only those bipartitions that are common to all input trees, making the resulting tree highly conservative since it consists only of universally present bipartitions. In contrast, the extended majority-rule consensus tree starts with the bipartitions included in the standard majority-rule consensus tree and then greedily adds the remaining bipartitions in descending order of frequency, provided they are compatible with all those already included.

This balanced nature of the majority-rule consensus has a theoretical foundation. The majority-rule consensus tree minimizes the mean bipartition distance to the input trees (Barthélemy and McMorris 1986), making it a median tree with respect to this distance. The bipartition distance between two trees, also known as the Robinson-Foulds (RF) distance, is defined as the cardinality of the symmetric difference of their sets of bipartitions: it counts the number of bipartitions that are present in one tree but not the other. The balance of the majority rule reflects an equal weighting of two types of errors: false positives, where bipartitions appear in the consensus tree but are absent from the input tree, and false negatives, where bipartitions occurring in the input tree are missing from the consensus tree. In contrast, the strict consensus tree minimizes a metric that assigns zero weight to false negatives, while the extended majority-rule consensus tree attempts to minimize a metric that assigns zero weight to false positives. Consequently, all branches in the strict consensus tree have 100% support with respect to the input trees. In the majority-rule consensus tree, branches have support above 50%. The extended majority-rule consensus tree contains all branches with greater than 50% support, as well as branches with less than 50% support, which decreases the average branch support compared to the majority-rule consensus tree.

While the majority-rule consensus method achieves a practically appealing balance and has theoretical optimality with respect to the RF distance, it has a significant limitation: while it is less conservative than the strict consensus, it still often produces a highly unresolved consensus tree close to a star tree topology (i.e., the tree with no internal branches/bipartitions and no phylogenetic information). This drawback becomes particularly apparent when the number of taxa is large and the phylogenetic signal in the data is low, because the number of possible bipartitions grows exponentially with the number of taxa, in contrast to the linear increase in the number of bipartitions in a single tree. This is problematic given the advent of large datasets in recent years. Furthermore, the presence of even a few “rogue” taxa whose positions in the inferred trees are unstable can significantly reduce the number of branch occurrences (Wilkinson 1996). Consequently, it is often the case that the majority of input trees share only a few branches/bipartitions. To address this problem, Wilkinson (1996) proposed the concept of reduced consensus trees, which improve resolution by removing rogue taxa. Related approaches often use the consensus tree itself as a diagnostic or objective function; for example, several methods identify taxa whose removal increases the resolution (Aberer et al. 2013) or the information content (Smith 2022) of the consensus.

A similar phenomenon has been observed in the context of Felsenstein’s bootstrap support. As the number of taxa increases, bootstrap support values tend to decrease, particularly for deep branches, due to the low probability of finding exactly the same branch from the reference tree in the bootstrap trees. Lemoine et al. (2018) addressed this issue by proposing the use of the transfer bootstrap expectation (TBE) as an alternative measure of branch support. This approach uses the transfer distance to evaluate the similarity between bipartitions, replacing the binary presence/absence indicator of standard bootstrap support with a gradual measure that reflects varying degrees of similarity. With TBE, the support is a measure of the correctness of the inferred branch. One assumes that some taxa around the inferred branch are misplaced and must be transferred from one side of the branch to the other in order to recover the genuine branch. The TBE support can be interpreted as the number of these taxa, while accounting for the number of taxa in the tree and the depth of the given branch. This interpretation differs greatly from the interpretation of FBP support, which is roughly the statistical confidence level that the inferred branch is entirely correct (assuming no reconstruction bias, long-branch attraction, or model miss-specification). For more details on FBP and TBE supports and their interpretation, refer to Lemoine et al. (2018) and Zaharias et al. (2023).

In this study, we take an analogous approach to address the limitations of the majority-rule consensus method. We propose replacing the bipartition distance with a finer-grained dissimilarity measure and computing a median tree with respect to that measure. Specifically, we consider three dissimilarity measures between trees: the scaled-transfer dissimilarity, the unscaled-transfer dissimilarity, and the quartet distance. Similar to the majority-rule consensus tree, our goal is to minimize the sum of false positives and false negatives with equal weights. However, instead of considering bipartitions, we consider quartet tree topologies, for example. This allows our consensus trees to capture more detailed structural aspects of the trees being compared. They contain more branches than the majority-rule consensus tree. These extra branches are supported by less than 50% of the input trees, but generally the average branch support remains high, especially the transfer support. In general, our consensuses are significantly more resolved than the majority-rule consensus tree but less resolved than the extended majority-rule consensus tree.

In the following sections, we first review the properties of the majority-rule consensus tree and provide a detailed description of our proposed dissimilarity measures. We then describe our algorithms for efficiently computing approximate median trees with respect to these measures. The performance of our methods is evaluated through extensive experiments on both simulated and real data. For simulated data, we consider both situations of summarizing the posterior trees in a Bayesian setting and bootstrap trees in a maximum-likelihood framework. Our methods are then applied to a benchmark that was recently proposed in the Bayesian framework, and a comparison is made between them and the CCD-MAP and other standard approaches used to summarize tree posteriors. Lastly, we apply our methods to large real phylogenetic datasets: the Ebola virus 2013-2016 epidemic in West Africa, the mammalian phylogeny, and an HIV dataset comprising over nine thousand taxa. These experiments demonstrate that our proposed approaches achieve greater resolution and informativity than the majority-rule consensus, while maintaining an appropriate balance with branch support, particularly in challenging cases with low phylogenetic signal.

### New Methods, Benchmarks and Data

#### Phylogenetic Trees

In this paper, we focus on the consensus of phylogenetic trees which are unrooted and unweighted (i.e., branch lengths are not considered). Although standard tree inference methods typically yield binary (i.e., fully resolved) trees, we make no assumption that the input trees from which the consensus is derived are binary. Every branch in a phylogenetic tree corresponds to a bipartition (or split) of the taxon set. Given a bipartition *b* = *A*|*Ā*of a set of taxa, we call each of *A* and *Ā*a *part* of the bipartition *b*. Additionally, we define the *depth* of a bipartition *b* = *A*|*Ā*to be min {|*A*|, | Ā |} that is denoted by depth(*b*) (e.g., the stemming branch of a cherry clade, which contains two taxa, has a depth of 2). Given a tree *T*, the set of bipartitions of *T* is denoted by *B*(*T*). A bipartition is said to be *internal* if no part is reduced to a single taxon. A bipartition in which one part contains a unique taxon is said to be *trivial* since such bipartition is found in all trees with the same taxon set. An unrooted binary tree has *n* − 3 internal bipartitions, where *n* is the number of taxa, and a non-binary tree has fewer. We use the notation *B*_int_(*T*) for the set of internal bipartitions included in the tree *T*, and *B*_tips_(*T*) for the set of trivial bipartitions. The *branch resolution*, or simply *resolution*, of a tree *T* is defined as |*B*_int_(*T*)|/(*n* − 3). The *quartet resolution* of a tree *T* is defined as the proportion of quartets in *T* which have resolved quartet topologies (which equals 1.0 when *T* is binary and 0.0 when *T* is a star tree).

#### Majority-Rule Consensus Tree and Bipartition Distance

The *majority-rule consensus tree* contains only those bipartitions present in more than 50% of the input tree collection. The majority-rule consensus tree has been widely used because of its intuitive definition and ease of computation. It is also known that the majority-rule consensus is a median tree with respect to the bipartition (or RF) distance (Barthélemy and McMorris 1986): for a given set of input trees *T* = {*T*_1_, …, *T*_*N*_}, the majority-rule consensus tree *T*_maj_(*T*) satisfies

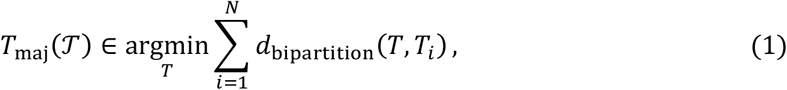

where *d*_bipartition_(*T, T*_*i*_) represents the *bipartition distance*. In short, the bipartition distance is the cardinality of the symmetric difference between two sets of bipartitions:

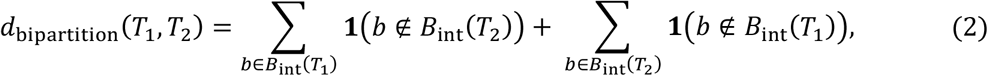

where **1**(*b* ∉ *B*(*T*)) is the indicator function that returns 1 when the branch *b* is not in *B*(*T*), and 0 otherwise. In the context of estimating the true tree *T*_2_ by an estimate *T*_1_, the first summation counts the number of the bipartitions in the estimate *T*_1_ but not in the true *T*_2_: it is the number of incorrect discoveries of bipartitions (i.e., false positives). The second term counts the number of the bipartitions in the true tree *T*_2_ that are not present in the estimate *T*_1_: it is the number of incorrect absences of bipartitions (i.e., false negatives). Thus, the bipartition distance gives equal costs to both false-positive and false-negative errors. The bipartition distance can be normalized by twice the maximum number of internal bipartitions, 2(*n* − 3), so that the normalized value will be in the range [0,1]. Note that if one of the trees (e.g., the consensus tree) is the star tree and the other (e.g., the true tree) is fully resolved, the normalized bipartition distance is 0.5. Thus, in this metric, any consensus having an error of 0.5 or more is worse than providing a star tree that essentially has no information (no non-trivial bipartitions).

As previously stated, a significant issue arises when the majority-rule consensus is applied to datasets comprising a large number of taxa and low signal. This can result in the generation of a highly unresolved consensus tree that closely resembles a star tree topology.

#### Quartet-based Fine-grained Dissimilarity Measure

The bipartition distance measures the number of branches that differ between two trees. In the case of a large number of taxa, it is possible for two trees to share many almost identical pairs of branches that are not exactly the same. This observation suggests that distances or dissimilarity measures capable of capturing finer gradations of similarity between bipartitions may lead to more accurate and meaningful consensus methods. This manuscript explores two types of measures that capture these similarities: the quartet-based approach and the transfer-distance-based approach.

The first approach is based on quartets, the sets of four taxa in the tree. When a set of four taxa (say *A, B, C, D*) is extracted from a tree, the quartet topology takes one of four possible forms: three resolved topologies (*AB*|*CD, AC*|*BD, AD*|*BC*) and one unresolved star topology. Let *QQ*(*T*) denote the set of all resolved quartet topologies observed in *T*. Then, the *symmetric quartet distance* is defined as follows:

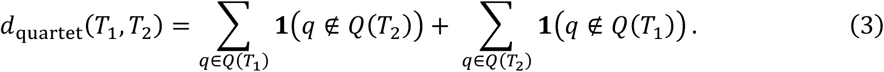

The definition is analogous to that of the bipartition distance, but in this case, the quartet topologies are employed in lieu of the bipartitions. In the context of estimating *T*_1_ by *T*_2_, this corresponds to equally weighting the false-positive resolved quartets (first summand) and false-negative resolved quartets (second summand), as with the bipartition distance (Eq. 2). By considering only the resolved quartet topologies in the summation, we distinguish two types of disagreement: when two trees have two different resolved topologies for a given quartet, the cost of that quartet is 2. On the other hand, when one of the trees has an unresolved quartet topology, and the other has a resolved one, the cost is 1. This type of quartet distance is sometimes referred to as the *parametric quartet distance* (Bansal et al. 2011), and this corresponds to the parameter *pp* = 0.5 case. For the sake of simplicity, this distance will henceforth be referred to as the *quartet distance*. The quartet distance can be normalized by dividing by 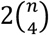, placing it in the range [0,1]. Similarly to the bipartition distance, when one of the trees is the star tree and the other is binary, the normalized quartet distance between them is 0.5. Thus, any consensus method that has more than 0.5 quartet distance to the true tree is worse than the star tree.

#### Transfer-based Fine-grained Dissimilarity Measures

The transfer distance is a metric distance between two bipartitions. Given two bipartitions *b*_1_ and *b*_2_, the transfer distance Transfer(*b*_1_, *b*_2_) is defined as the number of transfers required from one part of *b*_1_ to the other in order to match the bipartition *b*_2_ exactly. For example, consider the case where we have a set of eight taxa labeled *A* to *H*. The transfer distance between *b*_1_ = {*A, B, C, D*}|{*E, F, G, H*} and *b*_2_ = {*A, B, C, D, E, F*}|{*G, H*} is equal to 2, because one needs to transfer at least two taxa (*E* and *F*) from one side of *b*_1_ to the other to match the bipartition *b*_2_. Lin et al. (2012) defined the matching distance between binary trees based on the transfer distance between bipartitions. However, this distance is applicable only to binary trees, while our objective is to consider unresolved trees as well.

A novel branch support using the transfer distance was proposed by Lemoine et al. (2018). For a branch *b* in the reference tree *T* and the set of input trees 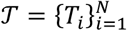, the transfer bootstrap expectation (TBE) is defined as follows:

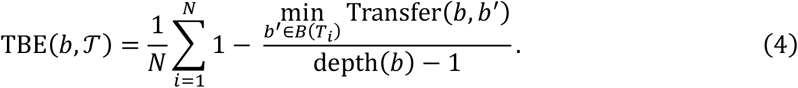

The minimum in the numerator of (4) is bounded above by depth(*b*) − 1. This is because for any bipartition *b* and any taxon *x* in the smaller part of the bipartition *b*, the transfer distance between *b* and the trivial bipartition {*x*} ∣ *XX* − {*x*}, where *XX* represents the set of all taxa, is depth(*b*) − 1. Thus, the quantity in the summand takes values in the range [0,1]. The same quantity can be computed in the case of Bayesian posterior tree draws when a reference tree is available. We refer to this support value as “transfer support” (TS), while the standard Felsenstein support value is called “frequency support” (FS) because it is equal to the frequency of the bipartition among the input trees.

We use a similar idea to define a dissimilarity measure between two trees as follows:

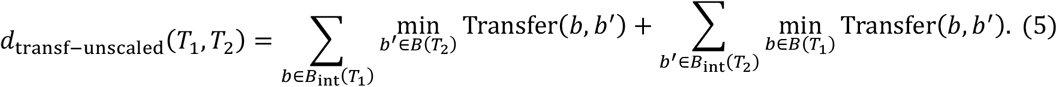

Each branch in *T*_1_ is penalized by the number of moves it needs to match one of the branches in *T*_2_, and vice versa. In the context of estimating *T*_1_ by *T*_2_, we again equally weight the false-positive errors (first summand) and false-negative ones (second summand). In this paper, we refer to this dissimilarity as *unscaled-transfer dissimilarity*. There are several possible ways to normalize this dissimilarity. One may divide the dissimilarity by the maximum number of internal branches 2(*n* − 3), in which case the normalized measure represents the average number of transfers it takes to match branches (although this will not be the case if either tree is unresolved). Another option is to divide by the maximum possible dissimilarity, which is 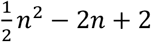 if *n* is even and 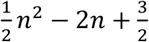 if *n* is odd (the proof is given in Supplementary Material). This maximum is attained when we have two very different caterpillar trees. This normalization results in a dissimilarity value falling within the range [0,1].

However, some deep branches can be very deep, which can result in heavy penalties. The number of transfers required can be as high as (depth(*b*) − 1) ~ *n*/2. Based on this observation, we also consider a scaled version of the transfer dissimilarity:

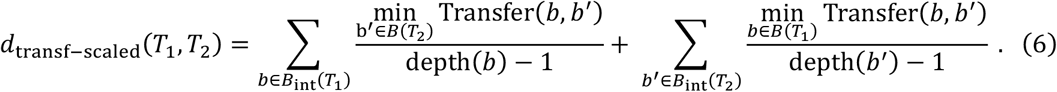

This scaling of each branch’s contribution to the dissimilarity ensures that each term within the sum takes values in [0,1]. This could be thought of as a graduation of the bipartition distance, where the exact presence/absence indicator is replaced by a gradual scale of dissimilarity. Thus, this scaled version of transfer dissimilarity is bounded above by the bipartition distance. Again, in the context of estimating *T*_1_ by *T*_2_, the false-positive errors (first summand) and false-negative errors (second summand) are given equal weight. We call this dissimilarity *scaled-transfer dissimilarity*. The whole measure can be normalized by dividing by 2(*n* − 3) to make the dissimilarity fall into the range [0,1]. Note that the scaled- and unscaled-transfer dissimilarities are, in general, not metric distances when *n* ≥ 6 (the triangle inequality is not guaranteed to hold).

The properties of these measures are summarized in Table 1. In the following, we call *unweighted* the bipartition (or RF) distance and scaled-transfer dissimilarity, since they give the same importance to every branch, regardless of its depth. We refer to the unscaled-transfer dissimilarity and quartet distance as *weighted*. Of all the dissimilarity measures considered here, the quartet distance gives the greatest weight to deep branches. From the bipartition viewpoint, a branch with parts of size *s* and *n* − *s* is compatible with 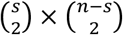 quartets, ranging from *O*(*n*^2^) for a shallow branch to *O*(*n*^4^) for a balanced deep branch. In this analysis, however, the same quartet may be counted for several bipartitions. From the quadripartition viewpoint, if the four subtrees incident to a branch contain *a, b, c*, and *d* taxa, that branch locally resolves *abcd* quartets. This number is *O*(*n*) for a shallow branch, can be *O*(*n*^2^) for a central branch in a caterpillar-like tree, and can reach *O*(*n*^4^) for a central branch in a balanced tree. In this analysis, some of the quartets are not counted, but the conclusion is the same: deep branches have more weight than shallow branches. Consequently, our quartet-based consensus method tends to favor deep branches, resolves the deep parts of the tree and significantly improves the quartet resolution, compared to the majority-rule consensus tree. The same holds true for unscaled-transfer dissimilarity, but to a lesser extent, since the range of branch errors is between 1 for a cherry and *∽n*/2 for a balanced deep bipartition.

**Table 1.** Properties of Dissimilarity Measures for Unrooted Phylogenetic Trees.

|  | Compares trees based on | Weight of branches |
| --- | --- | --- |
| **Bipartition (or RF) distance (Eq. (2))** | Exact matching of bipartitions | Equal weights for all branches |
| **Scaled-transfer dissimilarity (Eq. 6)** | Similarity of bipartitions<br>using transfer distance | Equal weights for all branches |
| **Unscaled-transfer dissimilarity (Eq. 5)** | Similarity of bipartitions<br>using transfer distance | Higher weights for deep branches |
| **Quartet distance (Eq. 3)** | Exact matching of quartets | Very high weights for deep branches |

#### Greedy Optimization to Obtain Approximate Median Trees

Recall that, for a set *T* of trees, the majority-rule consensus tree is a median tree of the trees in *T* with respect to the bipartition distance. For any fine-grained tree dissimilarity measure *d* among the three introduced above (i.e., the scaled- and unscaled-transfer dissimilarities, and the quartet distance), the consensus tree *C*_*d*_(*T*) considered in this paper is analogously a median tree of the trees in *T* with respect to *d*. In other words, *C*_*d*_(*T*) is a tree that minimizes the loss *L*_*d*_ defined by

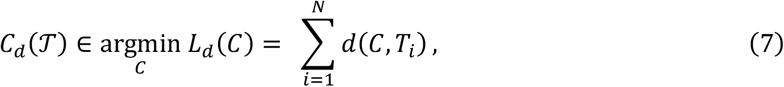

where *C* ranges over all possible tree topologies, both fully and non-fully resolved. Theoretical analyses (unicity of the solution, NP-hardness of the optimization task, …) are beyond the scope of this paper. Here, we propose and use greedy algorithms to obtain an approximation of the median tree *C*_*d*_(*T*). Importantly, our approach should not be confused with the extended majority-rule consensus tree, which is also based on a greedy algorithm. The latter seeks to add branches to the majority-rule consensus tree, with FBP support as high as possible (but less than 50%), which decreases false negative errors, but increases false positive errors (in terms of bipartitions). In our proposed consensus methods, both types of errors (in terms of quartet topologies and taxon transfers) are equally weighted, and our goal is comparable to the majority rule. We use greedy algorithms because no exact algorithms are known. We consider the following two algorithmic strategies:

##### Strategy 1

Prepare a fully resolved initial tree, repeatedly prune branches that reduce the loss *L*_*d*_ the most in a greedy manner, and stop when there are no more internal branches whose removal does reduce *L*_*d*_.

##### Strategy 2

First, prepare an initial consensus tree and a set of candidate branches. Then, greedily add or prune branches that reduce *L*_*d*_ the most. Stop when neither adding a candidate branch nor pruning a branch in the consensus tree can further reduce *L*_*d*_. Due to the exponential number of possible branches (bipartitions), we typically reduce the set of candidate branches to those that occur at least once in the input trees.

For the quartet-based median tree, we used simple solutions with existing software (tqDist; Sand et al. 2014) to efficiently compute quartet distances during each iteration of the approximation procedure (see Supplementary Material for details). To compute an approximate median tree with respect to transfer-based dissimilarities, we have developed an efficient algorithm based on the first pruning strategy above. To speed up the computations, we generalized a fast algorithm for the calculation of transfer supports (Truszkowski et al. 2019). In the remainder of this subsection, we provide an outline of this pruning algorithm. Further details can be found in the Supplementary Material.

At the core of the pruning algorithm are two procedures, which we now explain. Suppose we are given a fully resolved initial tree *T* with *n* taxa and a set of *N* input trees 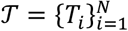 with the same set of taxa. In the first procedure, for each internal bipartition in the initial tree *T*, the algorithm computes its transfer support based on the set *T* of input trees. The transfer support can be calculated in *O*(*nN*(log *n*)^3^) time using the algorithm of Truszkowski et al. (2019). In the second procedure, for each bipartition in each input tree *T*_*i*_ in *T*, the algorithm computes its top *K* matching branches in the initial tree *T* by finding its closest *K* branches in terms of the transfer distance. *K* is a hyper-parameter of the method, which is much smaller than *n* (e.g., *K* = 30 with the HIV dataset where trees have more than 9,000 leaves). This computation can be done efficiently by generalizing the algorithm of Truszkowski et al. (2019) such that it finds top *K* bipartitions instead of just a single best one. Computing top *K* best matches for *O*(*nN*) bipartitions takes *O*(*nNK*(log *n*)^3^) time. If *K*≪*n*, this remains significantly faster than any naïve approach that requires explicit computation of all *O*(*n*^2^*N*) pairs of transfer distances.

The above two procedures are useful to quickly evaluate the benefit of pruning branches in the initial tree. For the unscaled-transfer dissimilarity, the loss *L*_*d*_ can be written as follows:

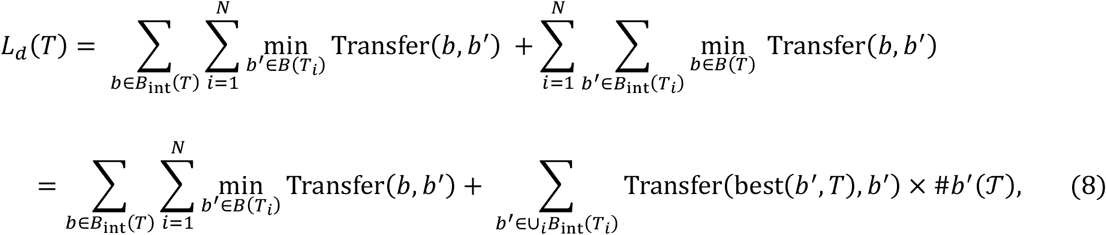

where best(*b*^′^, *T*) denotes one of the closest bipartitions to *b*^′^ in *T* (ties are broken arbitrarily), and #*b*^′^(*T*) is the number of appearances of the bipartition *b*^′^ in the set *T* of input trees. The first term of (8) represents the false-positive component of the loss and each transfer support value is calculated by the first procedure. If we prune a branch *b* of the initial tree *T*, the first term decreases accordingly. The second term of (8) represents the false-negative loss. This part can be rewritten as follows:

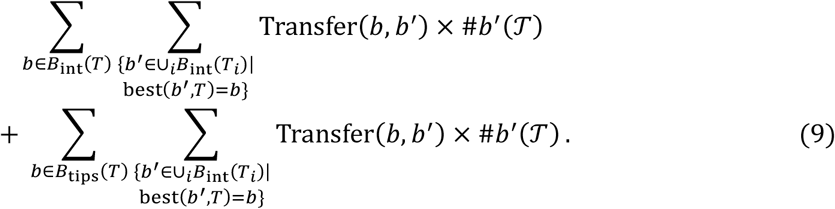

This form shows that if we prune an internal branch *b* from *T*, the false-negative loss increases by the following amount:

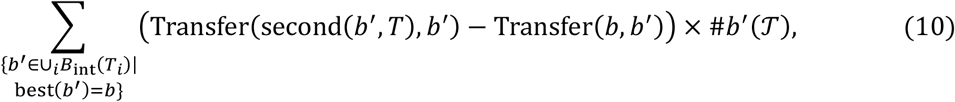

where second(*b*^′^, *T*) represents a second closest bipartition to *b*^′^ in *T*. Note that when several closest bipartitions exist, the difference in Eq. (10) becomes zero, and therefore the pruning benefit in Eq. (8) does not depend on the arbitrary tie breaking. Using the second procedure to find top *K* matches for all bipartitions of *T*, as well as their associated transfer distances, we can quickly compute the increase in the false-negative loss induced by the pruning of any given branch of *T*. Therefore, by iteratively updating the current “best match” and “second best match” for all internal bipartitions present in *T*, we can rapidly prune branches in *T* greedily based on loss reduction.

The purpose of computing the top *K* matches is to minimize the need for re-computation of the first and second-best matches. However, during the pruning process, the first or second matches of some branches may fall outside the original top *K* matches. When this happens, we use the algorithm of Lutteropp et al. (2020) to compute all transfer distances between the target branch in *T* and all branches in *T* in *O*(*n*) time and sort the distances in *O*(*n* log (*n*)) time. To efficiently store the numerous bipartitions induced by large collections of big trees together with their associated values (e.g., the top *K* matches), we use the hash table proposed by Amenta et al. (2003).

The total running time depends on the number of iterations and re-computations required during pruning. If only *O*(*NK*(*log n*)^2^) branches require re-computation, the total running time is as fast as *O*(*nNK*(*log n*)^3^), being almost linear with respect to the number of taxa *n* and the number of input trees *N*. In the worst case, where re-computation is performed for *O*(*nN*) bipartitions, the additional cost is *O*(*n*^2^*N* log (*n*)). In practice, for moderate *K* values, we are clearly far from the worst case. For the HIV dataset that will be introduced later, which has 9,147 taxa and 1,000 input trees, we use *K* = 30. The pruning computation took about 20 minutes on an ordinary laptop computer. (MacBook Pro, CPU: Apple M2 Pro, clock speed 3490 MHz, memory 32GB). This is typically much faster than computations required for the inference of the input trees, such as bootstrapping maximum-likelihood tree construction or Bayesian MCMC methods. The algorithm for the scaled-transfer dissimilarity is essentially the same, with similar computing times.

#### Simulated Data

The performance of each method was first evaluated using numerical simulations. Simulated datasets comprising 100 sequences (taxa) were generated as follows. First, 100 random trees with 100 leaves were generated using the Birth-Death model, with a birth rate of 0.2 and a death rate of 0.1. Following the general sampling approach proposed by Hartmann et al. (2010), the Birth-Death model was simulated until the tree contained 100×10 = 1,000 taxa. Then, a time when exactly 100 taxa were present was randomly selected, and the tree configuration at that time was extracted. This was done using the DendroPy package (Moreno et al. 2024). The height of each tree was normalized to either 0.05 or 0.5. These two tree heights represent relatively small and large evolutionary changes while remaining within a realistic range. After normalization, each branch length was multiplied by an independent lognormal random variable with a mean of 1.0 and a standard deviation of 0.5, thereby introducing a deviation from the unrealistic molecular clock hypothesis. Finally, to avoid generating an excessive number of identical sequence pairs, the minimum branch length was set to 0.5/*L*, where *L* is the sequence length. Specifically, all branch lengths below the specified threshold were replaced by either 0.5/100 or 0.5/300, depending on the sequence length. This ensured that the minimum number of expected mutations along each branch of the tree was 0.5. The final trees were obtained as unrooted trees by removing the root.

For each tree, two datasets of aligned sequences of lengths 100 and 300 were generated using Seq-Gen (Rambaut and Grass 1997). The HKY model was used as the substitution model and the nucleotide frequencies were set as: A=0.15, C=0.35, G=0.35 and T=0.15. The transition/ transversion ratio was set to 2.0 (corresponding to a transition rate 4 times faster than the transversion rate), and a gamma heterogeneity of rates across sites was introduced with parameter α = 0.5. These parameter values are all realistic and correspond to standard estimations found in real datasets.

#### Compared Methods and Criteria

We considered two approaches to analyze these datasets. The first approach is a Bayesian one, where we summarized the collection of trees obtained by sampling from the posterior distribution using an MCMC algorithm. The computation was done using MrBayes (Ronquist et al. 2012). The GTR+Γ model was used for the analyses as it is the most commonly used substitution model today. The MCMC chain was run for 1,200,000 generations, with the first 200,000 generations discarded as a burn-in period and the remainder sampled at a frequency of 1,000, generating a total of 1,000 tree draws from the posterior. Finally, the consensus of these 1,000 posterior trees was calculated using the different methods studied here.

The second approach combines maximum-likelihood estimation of trees with bootstrapping. RAxML-NG (Kozlov et al. 2019) was used with GTR+Γ model to estimate the maximum-likelihood tree as well as 1,000 non-parametric bootstrap trees. To stabilize the result, before entering the sequences into RAxML-NG, we extracted only one sequence from a set of identical sequences in the original dataset to make all input sequences unique. After analysis, the removed taxa were added back to form the unresolved clade of taxa with identical sequences. Finally, the consensus of the bootstrap trees was computed using the different methods studied here.

For the Bayesian approach, the following consensus trees were computed: majority rule, MCC (maximum clade credibility), MAP (maximum *a posteriori*), ASTRAL-IV corresponding to one of ASTER’s options (Zhang et al. 2025), EM (extended majority rule), and the three proposed consensus trees corresponding to the scaled-transfer dissimilarity, its unscaled version, and the quartet distance. The ASTRAL-IV algorithm seeks a binary tree that minimizes the false-negative component of the quartet distance loss. Since the false-negative and false-positive components of the quartet distance coincide in binary trees, ASTRAL-IV can be thought of as minimizing the quartet distance loss with the restriction that the output tree is binary. For the transfer approaches, the greedy addition and pruning algorithm (Strategy 2) was used, with the majority-rule consensus as the initial tree and the set of all bipartitions contained in the input tree collection as the candidate bipartition set. For the quartet approach, the ASTRAL-IV tree was used as the initial tree, and then the greedy pruning was applied (Strategy 1). This choice was based on the above-mentioned connection between the ASTRAL-IV tree and the symmetric quartet distance.

For the bootstrap analysis, the following (consensus) trees were considered: MLE (maximum-likelihood estimate) constructed by RAxML-NG, majority rule, EM, ASTRAL-IV, and the three consensus trees corresponding to the scaled-transfer dissimilarity, its unscaled version, and the quartet distance.

Each consensus tree was first evaluated by the mean dissimilarities to the input trees. We used the three dissimilarity measures considered above (the quartet distance, the scaled-transfer dissimilarity, and the unscaled-transfer dissimilarity) as well as the bipartition (RF) distance. These dissimilarity values show how well each consensus tree represents the input trees. The various consensus trees were also evaluated against the correct tree using the same set of dissimilarity measures. These dissimilarities indicate how well each consensus tree estimates the true topology used to generate the sequences. In addition to accuracy, the resolution of each consensus tree was evaluated using two measures: branch resolution and quartet resolution.

#### Bayesian benchmark

We evaluated our consensus methods on two simulated Bayesian phylogenetic datasets, Coal320 (320 taxa) and Yule400 (400 taxa), introduced by Berling et al. (2025). The objective of this experiment was to assess the performance of our consensus procedures in a Bayesian setting by reusing a recent benchmark and adopting the same evaluation criteria as in the original study. In Coal320, true trees were generated under a time-stamped coalescent model, and in Yule400 under a Yule pure-birth process. In both datasets, sequence alignments were simulated under an HKY+Γ substitution model, with alignment length 250 for Coal320 and 300 for Yule400. Each dataset comprises 100 independent replicates. For every replicate, the results of two independent BEAST2 MCMC analyses are provided, each consisting of 35,000 posterior trees.

For each replicate, we subsampled posterior trees from the two runs, reducing them to 500 trees per run, and then pooled them into a combined set of 1,000 trees. Each 1,000-tree set was used to construct the following consensus trees: majority rule, EM, CCD0-MAP (Berling et al. 2025), ASTRAL-IV, and the three proposed consensus trees based on scaled- and unscaled-transfer dissimilarities, and quartet distance, all obtained by pruning ASTRAL-IV tree. CCD1-MAP was excluded, as it was reported by Berling et al. (2025) to perform worse than CCD0-MAP on these datasets. All trees were treated as unrooted and branch lengths were ignored. The Coal320 and Yule400 posterior samples consist of rooted time trees. CCD0-MAP was computed in its native rooted setting using clades. For comparison with our methods, the root was subsequently removed from all consensus and reference trees, and only their unrooted topologies were evaluated. As in (Berling et al. 2025), we compared the consensus trees to the true trees using the RF (bipartition) distance. We also measured the resolution of the consensus trees and their fine-grained dissimilarities to the true tree, as in the simulation study described above.

To assess stability with respect to independent posterior sampling, we followed Berling et al. (2025) and evaluated the distance between consensus trees obtained from the two independent MCMC runs. For each replicate and each consensus method, we constructed consensus trees separately from each run using subsamples of size 3, 10, 30, 100, 300, and 1,000 trees per run, and computed the RF distance between the two resulting trees. The “precision” values obtained in this way were averaged across the 100 replicates for both Coal320 and Yule400. Smaller values indicate greater stability of the consensus method across independent posterior samples. This is an important aspect for detecting methods that tend to produce unstable and unreliable trees.

#### EBOV Data

We analyzed the dataset of Dudas et al. (2017) on the West African Ebola virus (EBOV) 2013-2016 epidemic, which comprises 1,610 complete genomes sampled between March 2014 and October 2015. The associated metadata describes how the virus spread and includes information about the country where the samples were taken, coded as Guinea, Sierra Leone, Liberia, or unknown. Baele et al. (2025) used this dataset to evaluate the performance of their HIPSTR consensus tree method in a Bayesian context. The original Bayesian analysis was performed in BEAST 1.8.4 using an HKY+Γ substitution model, a relaxed log-normal molecular clock, and a Skygrid coalescent prior. We randomly subsampled 1,000 trees from the 9,000 post-burn-in posterior trees. We computed the majority-rule consensus, EM, HIPSTR, and the three proposed consensus trees obtained by pruning from the ASTRAL tree (Strategy 1).

We evaluated the branch and quartet resolutions, as well as the average branch and transfer supports. We also compared the distributions of internal-branch depths with those of the input posterior trees. Since no reference tree is available, we used dendrogram purity (DP; Heller and Ghahramani 2005), adapted to unrooted trees, to evaluate the geographic coherence of the different consensus trees based on country labels. For each pair of tips from the same country, standard DP considers the cluster defined by their most recent common ancestor; in our unrooted setting, we instead use the smallest side of a bipartition containing both tips and calculate the fraction of tips in that cluster from the same country. When several minimum-size clusters exist, we use the one with the highest same-country purity. These values are averaged over same-country pairs and then across countries. Thus, DP values are high when tips with the same country label tend to form relatively pure clusters, and low when clusters contain a broad spectrum of labels. For example, if the consensus is under-resolved with large impure clusters, the DP value will be low. Conversely, a large pure cluster will increase DP, even if its topology is not fully correct. In cases of epidemic spread where multiple introductions are probable, we do not expect DP to be close to 1. However, a low DP value suggests that the country labels are poorly associated with the given consensus tree or that the resolution is low. Simple calculations show that, in the absence of phylogenetic association, the DP value is roughly equal to the inverse of the number of labels.

#### Mammal Data

To apply our methods to real-world data, we used the COI-5P mammalian amino acid sequence dataset from Lemoine et al. (2018). The original dataset contains 1,449 sequences from 527 sites, with the NCBI taxonomy (https://www.ncbi.nlm.nih.gov/taxonomy) serving as the reference tree for our analyses.

For the full dataset (1,449 taxa), we constructed four consensus trees directly from the bootstrap trees provided in Lemoine et al. (2018): majority rule, EM, and the two transfer-based consensus trees (scaled and unscaled), both obtained by pruning the MLE tree. The quartet-based consensus approach was not used with this large dataset for computing time reasons. While the NCBI taxonomy represents current knowledge of mammalian phylogeny, it is not a binary (i.e., fully resolved) tree. As a result, the bipartition distance and the fine-grained dissimilarity measures would overestimate the false-positive loss for highly resolved consensus trees. For quartets, restricting comparisons to the set of quartets resolved in the NCBI reference tree mitigates this issue. Therefore, in the full-dataset analysis, we mainly evaluated topological accuracy using the symmetric quartet difference, which was restricted to quartets resolved in the NCBI tree. The bipartition distance is also reported in the results for reference. Additionally, we assessed biological interpretability by examining the recovery of the nine mammalian clades reported in Lemoine et al. (2018). We considered a clade to be “approximately recovered” if its closest matching clade in the consensus tree had a transfer distance error of 15% or less. A clade was considered “fully recovered” if its closest matching clade was identical with a transfer distance of zero.

To enable a fair comparison using all dissimilarity measures, we further constructed subsampled datasets for which the NCBI taxonomy is fully binary. The largest subsets satisfying this requirement contain 164 taxa, and we randomly chose 100 such subsets. On average, the taxon overlap between two of these subsets is about 55%. This means that, although they are not independent, the 100 replicates provide complementary bases for method assessment. For each of these subsampled datasets, we estimated the maximum-likelihood tree and 1,000 maximum-likelihood bootstrap trees. RAxML-NG was used with mtMAM substitution model, which was specifically derived for analyzing mammalian mitochondrial protein sequences (Yang et al. 1998). A gamma model (+Γ) was used to account for the variability of rates across sites. We then computed consensus trees for each set of bootstrap trees using the same set of methods as in the simulation experiment: MLE, majority rule, EM, ASTRAL-IV, and the three consensus trees with respect to the scaled-transfer dissimilarity, the unscaled-transfer dissimilarity, and the quartet distance. Greedy addition and pruning from the majority-rule consensus tree (Strategy 2) was used for the transfer approaches, while pruning from the ASTRAL-IV tree (Strategy 1) was used for the quartet approach. We then compared each consensus tree to the fully-resolved NCBI reference subtree using each of the three proposed dissimilarity measures and the RF distance.

#### HIV Data

We further examined a previously published dataset (Lemoine et al. 2018) comprising an alignment of 9,147 HIV-1 *pol* gene sequences. The dataset contains *pol* sequences from the nine subtypes (A, B, C, D, F, G, H, J, and K) of the HIV-1 group M (HIV-1M). These sequences are annotated as non-recombinant (“pure” strains) in the Los Alamos HIV Sequence Database (https://www.hiv.lanl.gov/). However, among these sequences, Lemoine et al. (2018) identified 48 recombinants using the “jumping profile hidden Markov model” (jpHMM) software developed by Schultz et al. (2009). Each sequence was classified into one of the subtypes or into the recombinant category, based on jpHMM predictions. In this analysis, we used the reference and bootstrap MLE trees from Lemoine et al. (2018) to compute consensus trees.

A critical difference from the previous analysis of mammal data is that we do not have a reliable true tree for the HIV dataset, making it difficult to assess the goodness of consensus trees. Here, we first assess the mean dissimilarity to the input trees and the mean (transfer and frequency) branch support values to indicate how well each consensus tree summarizes the given tree collection. The mean support criterion tends to favor more unresolved trees, as including more branches will likely decrease the mean support, and vice versa. There needs to be a compromise between resolution and support. Furthermore, we evaluated how successfully each consensus tree recovers the nine subtypes present in the dataset. In this computation, recombinants were not considered. In a consensus tree, a given subtype is considered “fully recovered” if one of its subtrees contains all the sequences annotated with this subtype, possibly along with some recombinants, but no sequences annotated with another subtype. Since Lemoine et al. (2018) found that some subtypes had low Felsenstein bootstrap support, it is expected that the majority-rule consensus tree will fail to recover some of the subtypes. Thus, we searched the majority-rule consensus tree to check whether some of the subtypes were “approximately recovered”. For this purpose, we used the same transfer-based approach as in Lemoine et al. (2018), in which recombinants were not counted either.

Lastly, to gain additional insight into tree structure and the amount of phylogenetic information captured, we examined the overall resolution of each (consensus) tree and the depth of its bipartitions. These metrics were combined to assess whether the consensus trees are capable of adequately representing the phylogenetically significant groupings and maintains crucial information about HIV subtypes, while showing a reasonable level of resolution, given the low signal in this data and the huge number of sequences (tree leaves).

For computational reasons, we focused on the following set of (consensus) trees: MLE, majority rule, EM, ASTRAL-IV, and the two transfer-based consensuses obtained by pruning the MLE tree (Strategy 1).

#### PhyloCRISP Software

We provide an implementation of the proposed methods in the PhyloCRISP software package. PhyloCRISP (Phylogenetic Consensus Resolution Improvement using Split Proximities) implements both optimization strategies for transfer-based dissimilarities (scaled and unscaled) as well as the pruning-based strategy for median-tree construction under quartet distance. PhyloCRISP is available at https://github.com/yukiregista/PhyloCRISP. It outputs the resulting consensus trees with both frequency support and transfer support values. Detailed instructions for installation, usage, and input/output formats are provided in the Supplementary Material, along with a computing time comparison with other consensus software using a medium-sized dataset, Yule400 (400 taxa; see above). Using this dataset, all of our consensus algorithms take only a few dozen seconds on a laptop (Supplementary Table S11), which is considerably less time than it takes to infer the collection of input trees using a Bayesian or bootstrap approach. The quartet-based consensus becomes substantially slower with larger datasets, while the transfer-based consensuses remain fairly fast.

## Results

### Bayesian Analysis of Simulated Data

Figure 1a shows the average dissimilarity between consensus trees and the input posterior trees in four different Bayesian settings, relative to the performance of the majority-rule consensus tree. The results for majority rule are provided in the Supplementary Material (Supplementary Tables S1 and S2). These results show how successful each consensus tree is in summarizing the distribution of input trees, as gauged by four dissimilarity measures. The four median trees considered here (the majority rule and the three proposed consensus trees) attempt to minimize the sum of the four studied dissimilarity measures with respect to the input trees. The results show that the optimizations are successful to the extent that each computed consensus tree is the best in terms of its corresponding dissimilarity measure. The majority rule is the best with respect to the bipartition (RF) distance, as expected. However, proposed consensus approaches show only a small increase (up to 5%) in the bipartition loss compared to the majority rule. Furthermore, the proposed approaches effectively summarize the input trees in terms of the other fine-grained dissimilarity measures, in addition to the ones being optimized. In fact, the three proposed approaches have comparable average dissimilarities to the input trees with respect to the three fine-grained metrics. One could hypothesize that this phenomenon is due to the resemblance of these dissimilarity measures. However, this cannot be the only explanation, as the performance of other methods varies significantly with the dissimilarities used. For example, the majority rule performs relatively well in terms of scaled-transfer dissimilarity, but poorly in terms of quartet distance. In contrast, ASTRAL-IV performs poorly with respect to the RF distance, but relatively well in terms of quartet distance, even though the inferred tree is fully resolved.

**Figure 1:**
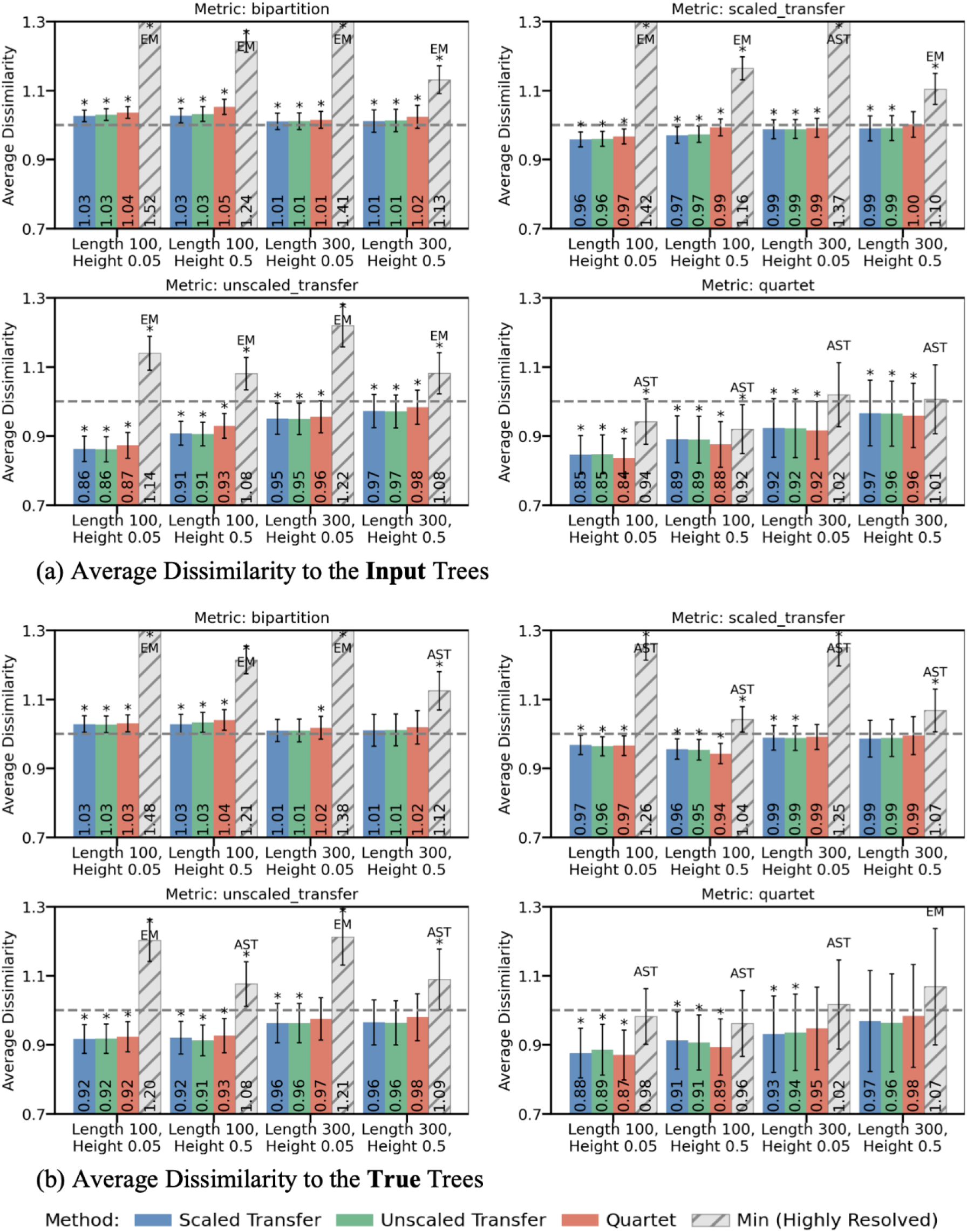
Simulated data, Bayesian setting. We compare the performance of our consensus trees to that of the majority-rule tree. For each proposed method, we show the ratio of its average dissimilarity to the target trees compared to that of the majority-rule consensus (dashed horizontal line at Y = 1.0). Two types of target trees are considered: **(a)** average dissimilarities to the input trees across 100 datasets; **(b)** average dissimilarities to the true trees across 100 datasets. The grey bar shows the minimum dissimilarity among four highly resolved approaches (EM: Extended Majority, AST: ASTRAL-IV, MCC, MAP), with the best-performing method labeled above each bar. The error bars indicate 95% confidence intervals, and asterisks indicate p-values (p<0.01) based on paired t-tests of differences between each estimator and the majority-rule consensus tree.

The reduction in average dissimilarities is particularly large for low signal, while it is small for high signal. In the lowest signal setting (sequence length = 100, tree height = 0.05), reductions of up to 4% in the scaled-transfer dissimilarity, up to 14% in the unscaled-transfer dissimilarity, and up to 16% in the quartet distance are observed. In the highest signal setting (sequence length = 300, tree height = 0.5), reductions of up to 1% in the scaled-transfer dissimilarity, up to 3% in the unscaled-transfer dissimilarity, and up to 4% in the quartet distance are observed.

In all settings, the highly resolved approaches (MAP, MCC, ASTRAL-IV, and EM) underperform the proposed approaches. This result is intuitive, as the dissimilarity measures considered in this paper attempt to balance the number of false positives and the number of false negatives, with varying definitions of false-positive and false-negative components. The generation of a highly resolved tree results in an excess of false positives, which in turn increases the average dissimilarity to the input trees, especially in cases where the signal is low. Furthermore, the highly resolved approaches are outperformed by the majority-rule approach, with the exception of the quartet distance, where ASTRAL-IV performs better than the majority rule in two out of four settings. This result can be attributed to ASTRAL-IV’s minimization of the quartet distance among all binary trees, as well as the discrepancy between the nature of the bipartition distance (minimized by the majority rule) and that of the quartet distance. In particular, as discussed earlier, deep branches are given significantly higher weight when using the quartet distance, while the majority rule treats each branch equally. Across all settings and for all four dissimilarity measures, the best highly resolved approaches in terms of average dissimilarity to the input trees are either ASTRAL-IV or EM. In contrast, the commonly used point estimates in Bayesian analysis, MAP and MCC, are consistently inferior. Berling et al. (2025) provide further results along these lines, as well as our subsequent results from our analysis of their Bayesian benchmark. See also the Supplementary Material (Supplementary Fig. S1) for a detailed comparison among highly resolved approaches.

Figure 1b shows the average dissimilarities between consensus trees and the true phylogeny. These results measure the performance of each consensus tree as a point estimator of the true phylogeny. The overall trend is similar to that described above for average dissimilarity against input trees. For all three fine-grained dissimilarity measures, the proposed approaches outperform the majority-rule approach. Furthermore, the loss reduction is higher in the low-signal cases, and the observations with highly resolved trees hold again. The similarity of the results is expected, as there are no model violations in this experiment (the substitution model GTR+Γ, which was used for tree inference, is more general than HKY+Γ, which was used for sequence simulation). In other words, we expect the median of the posterior distribution to be close to the true tree, with missing branches in the low-signal cases.

Figure 2a shows the resolution of the consensus trees in the four different settings. Overall, as shown in Figure 1, there are few differences between the three proposed approaches. The quartet-based consensus is slightly more resolved than the two transfer-based consensuses, which are almost identical. The proposed approaches demonstrate consistent improvements in both branch and quartet resolutions compared to the majority rule. For the lower signal cases with a sequence length of 100, the gain in resolution is higher, with the proposed approaches improving the branch resolution by 6-11% and the quartet resolution by 9-16%. In contrast, for the highest signal case, the gain in resolution is at most 4% for both types of resolution. The increase in resolution is distributed across branch-depth classes, rather than being concentrated at the shallowest or deepest branches, indicating that there is no evident depth-related bias (Supplementary Fig. S3a). The increased resolution is accompanied by only a modest reduction in average support relative to the majority-rule consensus (frequency support: 0.79–0.92 versus 0.86–0.93; transfer support: 0.89–0.96 versus 0.92–0.96; Supplementary Table S7). In fact, the average transfer support remains almost the same, demonstrating that transfer-based consensuses provide branches with high levels of transfer support, as expected. At the same time, the proposed consensus trees have substantially higher support than the highly resolved alternatives (frequency support: 0.42–0.85; transfer support: 0.68–0.92; Supplementary Table S7), particularly in the low-signal settings.

**Figure 2:**
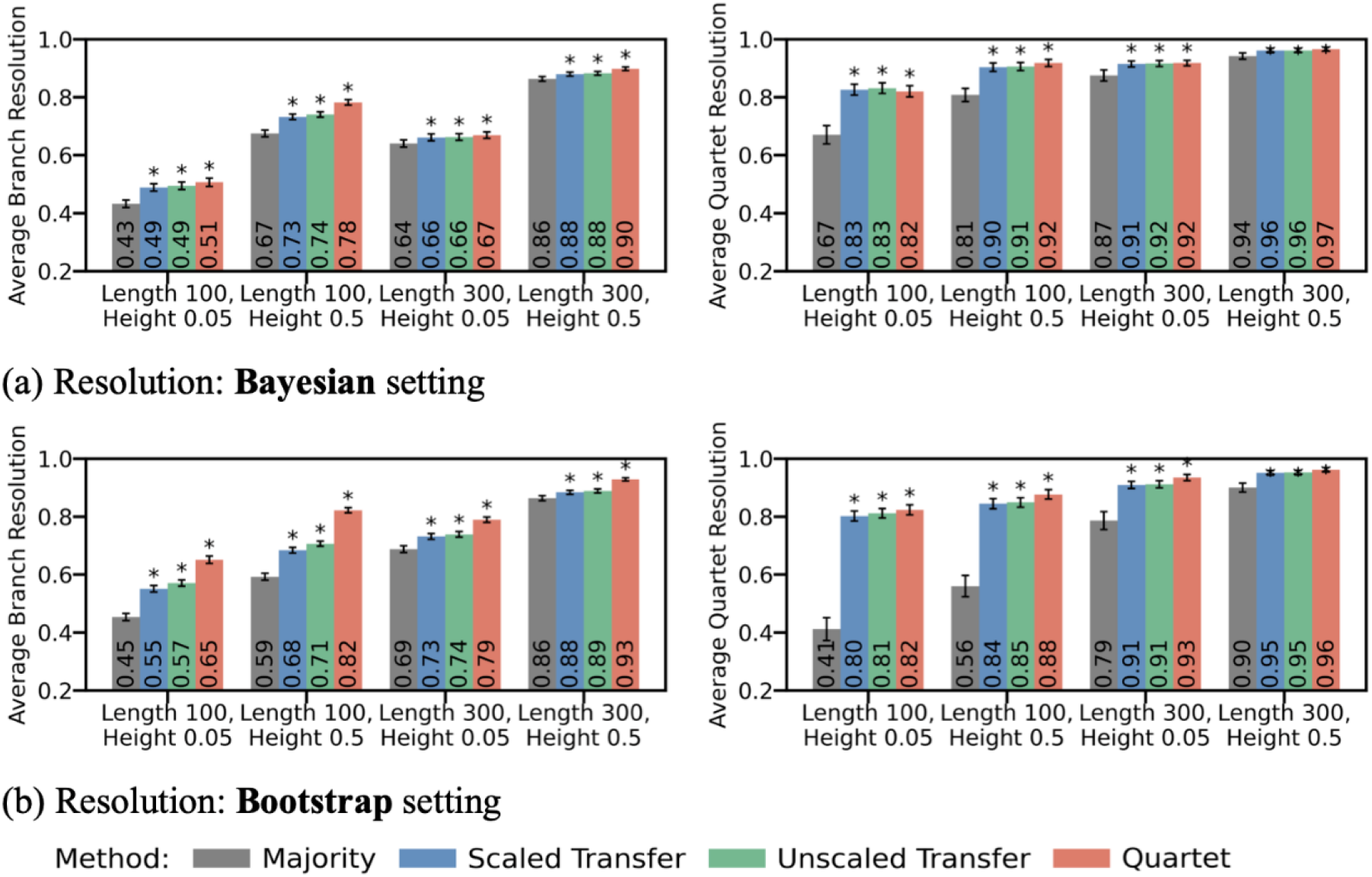
Simulated data, resolution of consensus trees. We plot the average branch resolution (left panels) and average quartet resolution (right panels) of consensus trees estimated by each method, across four simulation conditions defined by sequence length and tree height, for **(a)** the Bayesian setting and **(b)** the bootstrap setting. Both resolution measures are proportions ranging from 0 to 1, with 1 (100%) corresponding to full resolution. The error bars correspond to 95% confidence intervals of the average resolution of each estimator. The asterisks indicate p-values (p<0.01) of paired t-tests of resolution differences between each estimator and the majority-rule consensus tree.

#### Bootstrap Analysis of Simulated Data

Figure 3(a) shows the average dissimilarities between the consensus trees and the input trees, and Figure 3(b) shows the average dissimilarities to the true phylogeny, both relative to the majority-rule consensus tree. The results for the majority-rule consensus are provided in Supplementary Material (Supplementary Tables S3 and S4). The global tendency is consistent with the Bayesian case: the three proposed approaches are effective for all three dissimilarity measures, and the benefits of considering these approaches are more pronounced in low-signal settings. However, the degree of improvement is much more pronounced than in the Bayesian analyses. For example, in terms of the average dissimilarity between the consensus trees and the true tree, up to 18%, 36%, and 45% loss reduction (vs 6%, 9%, and 13% in the Bayesian setting) are observed for the scaled-transfer, unscaled-transfer, and quartet dissimilarities, respectively. Regarding resolution, the global trend is once again the same with the gain distributed across branch-depth classes (Supplementary Fig. S3b); however, the improvement is greater. For example, the quartet resolution of the majority-rule consensus in the lowest signal case is much lower (41%) than in the Bayesian setting (67%), but that of the proposed consensus trees is similar at around 80% (Fig. 2). The significant improvement in both loss reduction and resolution can be attributed to the high variance of the bootstrap distribution, which has a significant impact on the resolution of the majority-rule consensus tree. Overall, there are again few differences between the three proposed approaches. The quartet-based consensus is more resolved than the two transfer-based consensuses, but the accuracy in reconstructing the true tree is almost the same for all three approaches.

**Figure 3:**
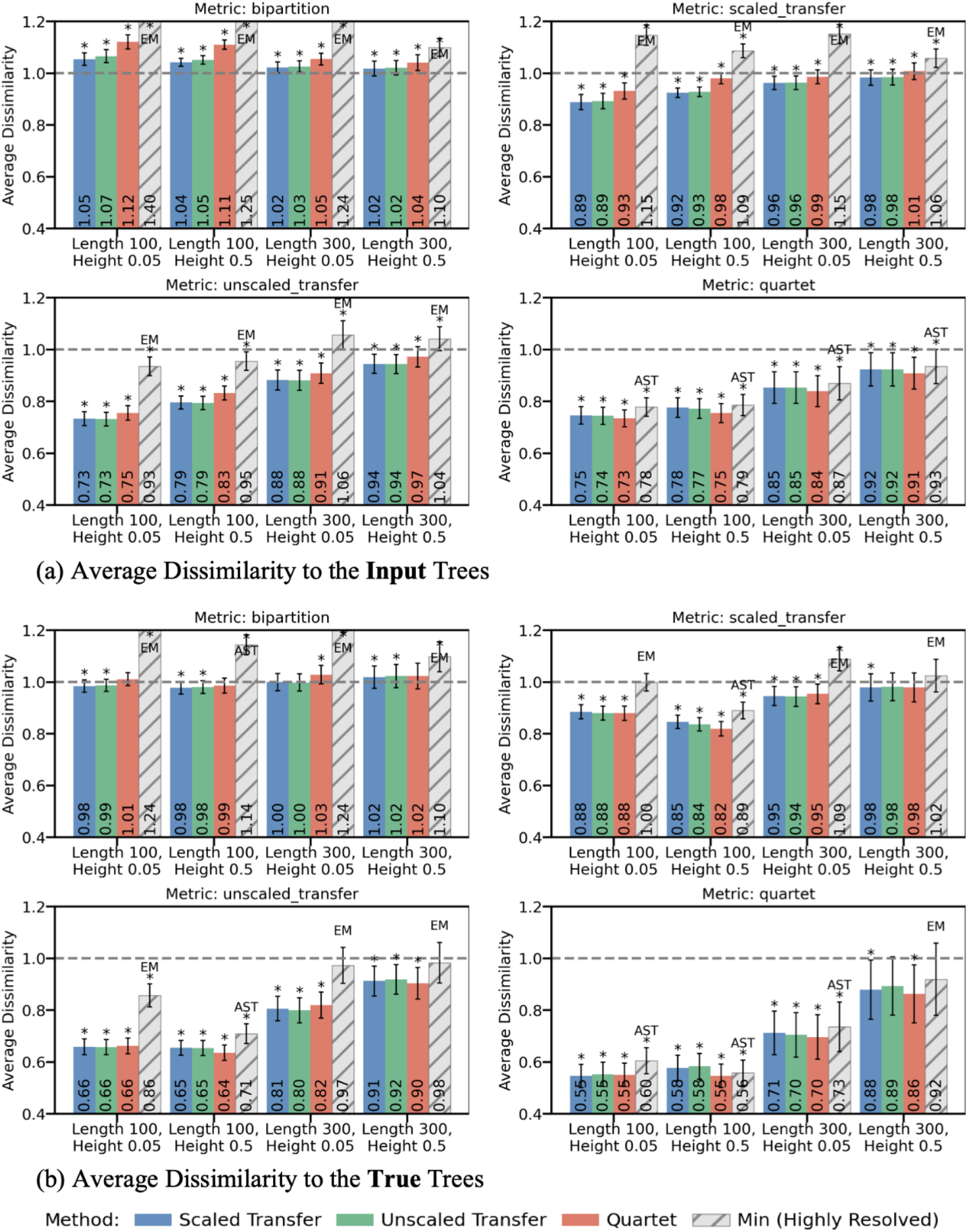
Simulated data, bootstrap setting. We compare the performance of our consensus trees to that of the majority-rule tree. For each proposed method, we show the ratio of its average dissimilarity to the target trees compared to that of the majority-rule consensus (dashed horizontal line at Y = 1.0). Two types of target trees are considered: **(a)** average dissimilarities to the input trees across 100 datasets; **(b)** average dissimilarities to the true trees across 100 datasets. The grey bar shows the minimum dissimilarity among highly resolved approaches (EM, AST: ASTRAL-IV, MLE), with the best-performing method labeled above each bar. The error bars are 95% confidence intervals, and asterisks indicate p-values (p<0.01) based on paired t-tests of differences between each estimator and the majority-rule consensus tree.

Highly resolved approaches tend to outperform the majority rule with respect to the weighted dissimilarities (unscaled-transfer and quartet-based), particularly in low-signal settings, whereas they tend to exhibit inferior performance for the unweighted measures (scaled-transfer and RF). The low resolution of the majority-rule consensus tree indicates that in the bootstrap case, the exact same deep branches do not appear as often as in the Bayesian case. This results in the majority rule being less accurate, especially in the weighted measures, due to a high false-negative rate that discards deep branches. In contrast, our approaches are able to detect the similarities of the deep branches and show a resolution comparable to the Bayesian case. Consistent with the larger gain in resolution, the proposed trees had lower average FBP support than the majority-rule consensus (0.66–0.86 versus 0.80–0.88; Table S8), while retaining similar TBE support values (0.81–0.93 versus 0.87– 0.93). Both support measures nevertheless remained consistently higher than those of the highly resolved alternatives (FBP: 0.47–0.81; TBE: 0.70–0.90).

Just as MAP and MCC never dominated ASTRAL-IV and EM, MLE is never the best approach among highly resolved ones (MLE, ASTRAL-IV, and EM). Although this may seem counterintuitive, it can be understood as reflecting the fact that these consensus approaches use more information provided by the set of input trees to assess the robustness and credibility of each inferred branch. This phenomenon of improving estimates by aggregating bootstrap estimates (“bagging”) is well known in the machine learning literature (Breiman 1996). See Supplementary Material (Supplementary Fig. S2) for detailed comparison among highly resolved approaches.

#### Bayesian benchmark analysis

In this section, we report the results of the Bayesian benchmark experiments on the Coal320 and Yule400 datasets, each containing 100 replicates and trees with 320 and 400 leaves, respectively. We compared majority rule, EM, our three proposed consensus methods, CCD0-MAP (best method from Berling et al., 2025), ASTRAL-IV, and MCC, which is the most widely used approach in a Bayesian setting. As in Berling et al. (2025), we considered two types of metrics. Firstly, the topological accuracy of the true tree reconstruction was measured using the RF distance and the three fine-grained dissimilarities. Secondly, we measured the stability of the various methods by comparing the consensus trees obtained from two different MCMC runs. The results are shown in Figure 4.

**Figure 4.**
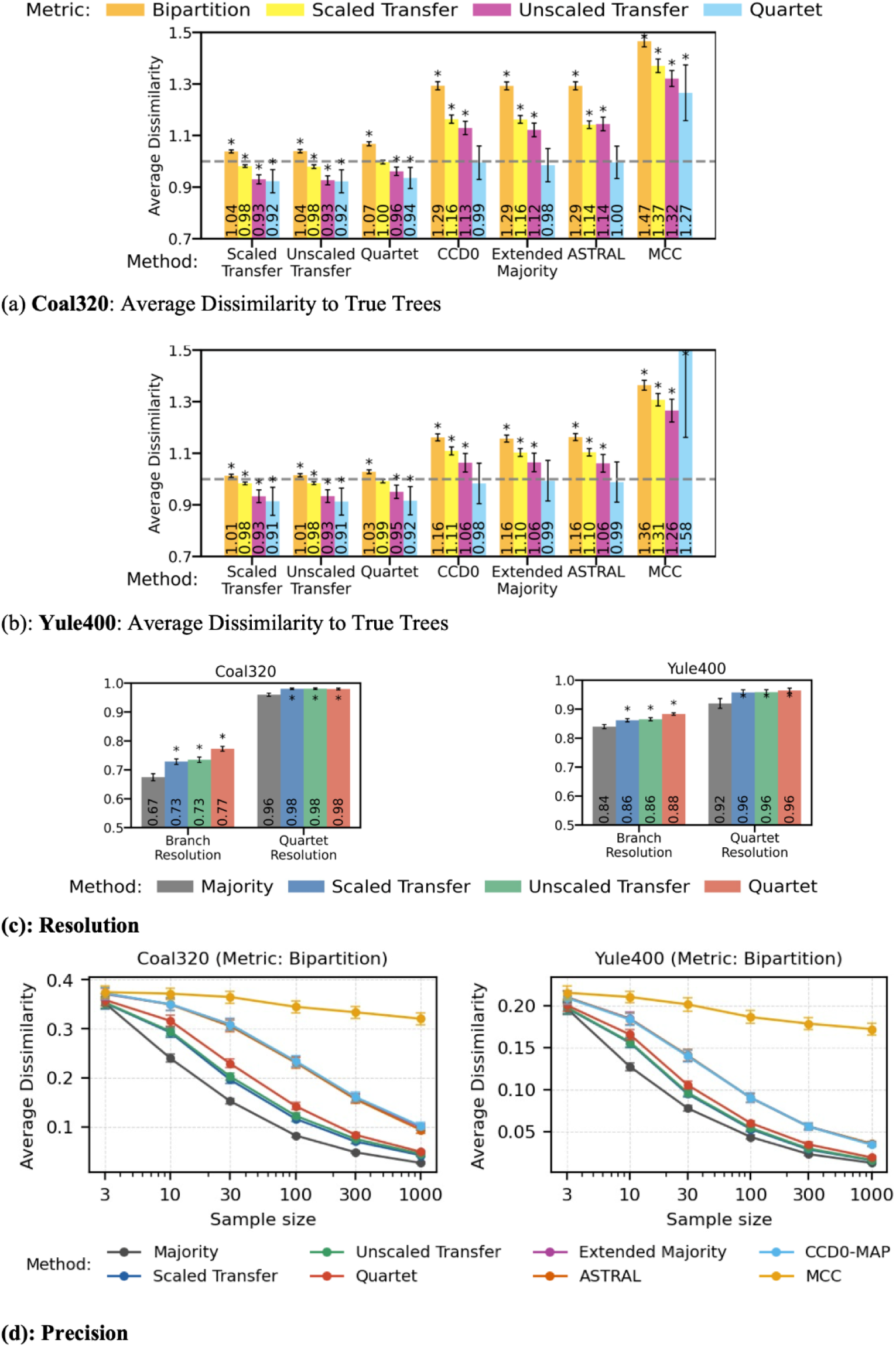
Bayesian benchmark. **(a)** Coal320 and **(b)** Yule400: average dissimilarities to the true trees, represented as a ratio of the estimator’s dissimilarity to the majority-rule’s dissimilarity (dashed horizontal line, Y=1.0). **(c)** Average branch resolution (BR) and quartet resolution (QR) for Coal320 (left) and Yule400 (right). **(d)** Precision analysis: average normalized bipartition distance between consensus trees obtained from two independent posterior subsamples, shown as a function of posterior sample size (left: Coal320 right: Yule400). Error bars are 95% confidence intervals. Asterisks indicate p-values (p < 0.01) from paired t-tests comparing each method to the majority-rule consensus tree.

The main message conveyed by Figure 4 (a-b) is that our three consensus methods and the majority rule provide superior topological accuracy compared to highly resolved consensus trees (CCD0-MAP, ASTRAL-IV, and EM). Figure 4 (a-b) also shows that MCC’s topological accuracy is significantly inferior to that of all other approaches. Moreover, there are only minor differences among our three consensus approaches. The quartet-based consensus is slightly more resolved than the other two approaches, which are very similar.

A consistent pattern emerges from the results of our three approaches across both datasets: a small increase in RF distance is accompanied by a reduction in fine-grained metrics compared to the majority-rule consensus. In Coal320, the reduction is relatively small. In Yule400, the RF distance remains similar to that of the majority-rule consensus (an increase of less than 3%), while the quartet distance to the true tree decreases by almost 10%. These improvements in fine-grained topological accuracy are accompanied by clear gains in resolution (Fig. 4c). The branch resolution increases from 0.67 under the majority-rule consensus to 0.73 for the transfer-based consensuses and to 0.77 for the quartet-based approach in the Coal320 dataset, with no depth-related bias (Supplementary Fig. S4). The proposed consensus trees retain high support, with transfer support nearly identical to that of the majority-rule consensus (0.94–0.96 versus 0.95–0.96) and substantially higher than that of EM, CCD0-MAP, ASTRAL-IV, and MCC (0.85–0.91).

The highly resolved consensus approaches (CCD0-MAP, EM, and ASTRAL-IV) display a broadly similar performance profile in Fig 4a-b. As expected, all three have substantially larger bipartition distances than majority rule, with increases of ~30% in Coal320 and ~15% in Yule400. They also show increased scaled- and unscaled-transfer dissimilarities compared with majority rule. These methods achieve slight improvements in quartet-based topological accuracy relative to majority rule (1-2%), but this gain is smaller in magnitude than that achieved by the proposed consensus methods (6-9%). In contrast, MCC performs distinctly worse, showing the largest increase across all dissimilarity measures (27% to 58%). The precision analysis (Fig. 4d) reveals the anticipated trade-off in stability between highly resolved trees and more conservative, partially resolved consensus trees. Consistent with its conservative nature, majority rule remains the most stable method across posterior sample sizes. The proposed consensus trees are markedly more stable than the highly resolved methods and rapidly converge as the posterior sample size increases. Among the proposed approaches, quartet-based consensus is less stable than transfer-based consensus, reflecting its higher resolution. Consistent with their dissimilarity profiles (Fig. 4a-b), CCD0-MAP, EM, and ASTRAL-IV exhibit nearly identical stability patterns across sample sizes and across both datasets. The MCC tree’s stability is substantially poorer than that of all other approaches.

#### Analysis of EBOV Data

Figure 5 summarizes the results for the West-African Ebola virus dataset, which includes 1,610 complete genomes associated with sampling-country annotations, and 1,000 posterior tree draws. The majority-rule consensus is conservative: it has high average support (FS = 0.90, TS = 0.94), but low resolution (BR = 0.43, QR = 0.57). The proposed consensus trees recover substantially more structure, increasing branch resolution to 0.52-0.57 and quartet resolution to 0.84-0.88. The gain is especially clear for quartet resolution, suggesting that the added branches capture broader tree structure rather than only introducing small shallow clades. This increased resolution was not accompanied by a large reduction in average support. The proposed trees retain high transfer support (TS = 0.89-0.92) and moderate-to-high frequency support (FS = 0.74-0.79). By contrast, HIPSTR and EM are nearly or fully resolved (BR = 0.98-1.00; QR = 0.91-1.00), but their average supports are much lower (FS = 0.48-0.49; TS = 0.71-0.72). Thus, the proposed methods occupy an intermediate regime: they recover much more resolution than majority rule while avoiding the weakly supported over-resolution seen in highly resolved alternatives. The branch-depth distribution profiles further show that the proposed consensus trees add branches across all depth bins relative to majority rule, without producing a pronounced excess at any particular depth.

**Figure 5.**
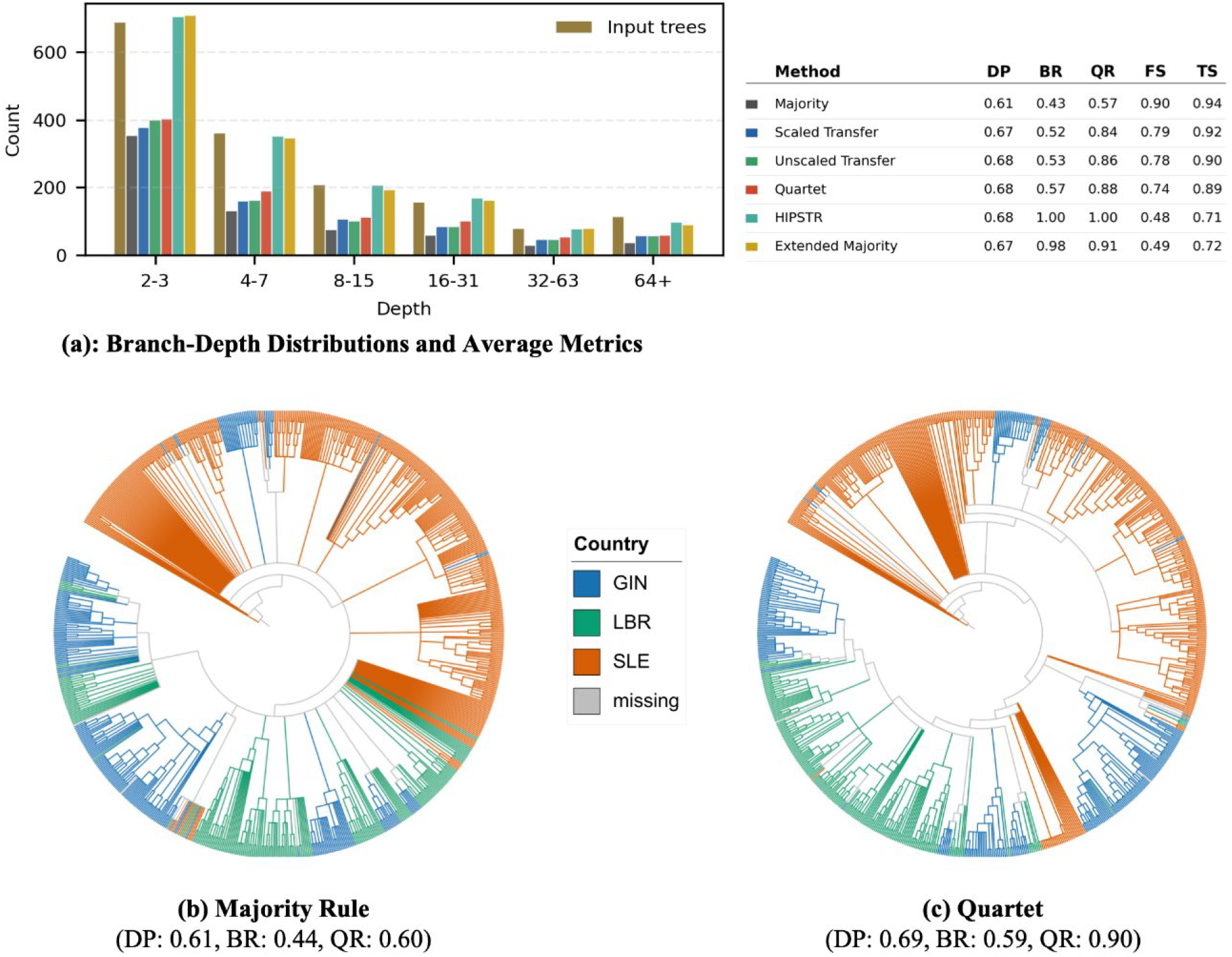
Comparison of consensus trees on the EBOV dataset. **(a)** Distribution of branch depths (left) and summary statistics (right) for each consensus method; DP: dendrogram purity; BR: branch resolution; QR: quartet resolution; FS: average frequency support; TS: average transfer support. **(b)** Majority-rule consensus tree. **(c)** Quartet-based consensus tree. Both trees are based on EBOV strains sampled on or before December 31, 2014 (810 tips). Branches are colored by the dominant sampling country among descendant tips when at least 90% of descendant labeled tips are from the same country; otherwise, branches are shown in gray.

Geographic coherence, measured by unrooted dendrogram purity (DP) over country labels, also improves. The majority rule obtains a DP value of 0.61, whereas the proposed methods reach values between 0.67 and 0.68. EM and HIPSTR achieve similar DP values (0.67 and 0.68, respectively). Thus, the proposed consensus trees nearly match the performance of highly resolved methods in recovering country-associated structure, while retaining substantially stronger support. Note that all these DP values are significantly larger than 1/3 (i.e., the inverse of the number of countries), indicating a clear association between the country labels and the given consensus tree.

Panels 5b and 5c compare the majority-rule and quartet consensus trees for the 810 samples collected on or before December 31, 2014. This subset highlights the same pattern: the majority-rule consensus remains relatively sparse, whereas the quartet consensus tree recovers additional internal structure, revealing more country-coherent groupings, which increase the DP value.

#### Analysis of Mammal Data

Results for the full mammal dataset (1,449 taxa, 1,000 bootstrap trees) are shown in Figure 6. Since the NCBI taxonomy is not fully resolved, topological accuracy was evaluated using the symmetric quartet difference, restricted to quartets that are resolved in the NCBI taxonomy. Compared to the majority-rule consensus tree, the transfer-based consensus trees substantially reduce the quartet distance (0.31 versus 0.46) while showing similar RF distance to the NCBI taxonomy (0.20-0.23 vs. 0.16). Majority rule is highly unresolved (branch resolution: 0.08, quartet resolution: 0.07) and approximately recovers only four of the nine considered mammalian clades, under the 15% transfer-error criterion. Transfer-based consensus trees achieve much higher branch (0.19–0.26) and quartet (0.62–0.63) resolutions. There are substantial differences between the two approaches (as opposed to Figs 1–5), with unscaled-transfer being more resolved than scaled-transfer. Both transfer-based consensuses approximately recover all nine clades. For example, the Mustelinae (weasels, ferrets, minks…) are fully recovered, whereas majority rule does not recover them. The Eulipotyphla clade (“true insectivores”, including hedgehogs, moles, and true shrews), which majority rule does not recover, is approximately recovered with 2 errors over 56 taxa (i.e., ~3.5%). The branch-depth distribution profiles (Fig. 6a) demonstrate once more that the proposed consensus trees introduce branches across all depth bins without exhibiting a significant excess at any specific depth. EM does not produce a fully resolved tree and has low quartet resolution (0.12), meaning it contains few deep branches. The branch-depth distribution of EM shows substantially more branches than the input trees at shallow depths, suggesting a bias toward including shallow branches. While EM also approximately recovers all nine clades, its quartet distance to the NCBI tree is higher (0.44) than that of transfer-based consensus trees (0.31). As expected, the MLE tree is far from the NCBI tree in terms of RF (bipartition) distance (0.59), although it is the same as our consensus trees for the quartet distance (0.31). Overall, the higher resolution of the transfer-based consensus trees is accompanied by lower average support than for majority rule (FBP: 0.37–0.43 versus 0.69; TBE: 0.56–0.64 versus 0.75), but their support remains significantly higher than that of EM, ASTRAL-IV, and MLE (FBP: 0.12–0.18; TBE: 0.33–0.35).

**Figure 6.**
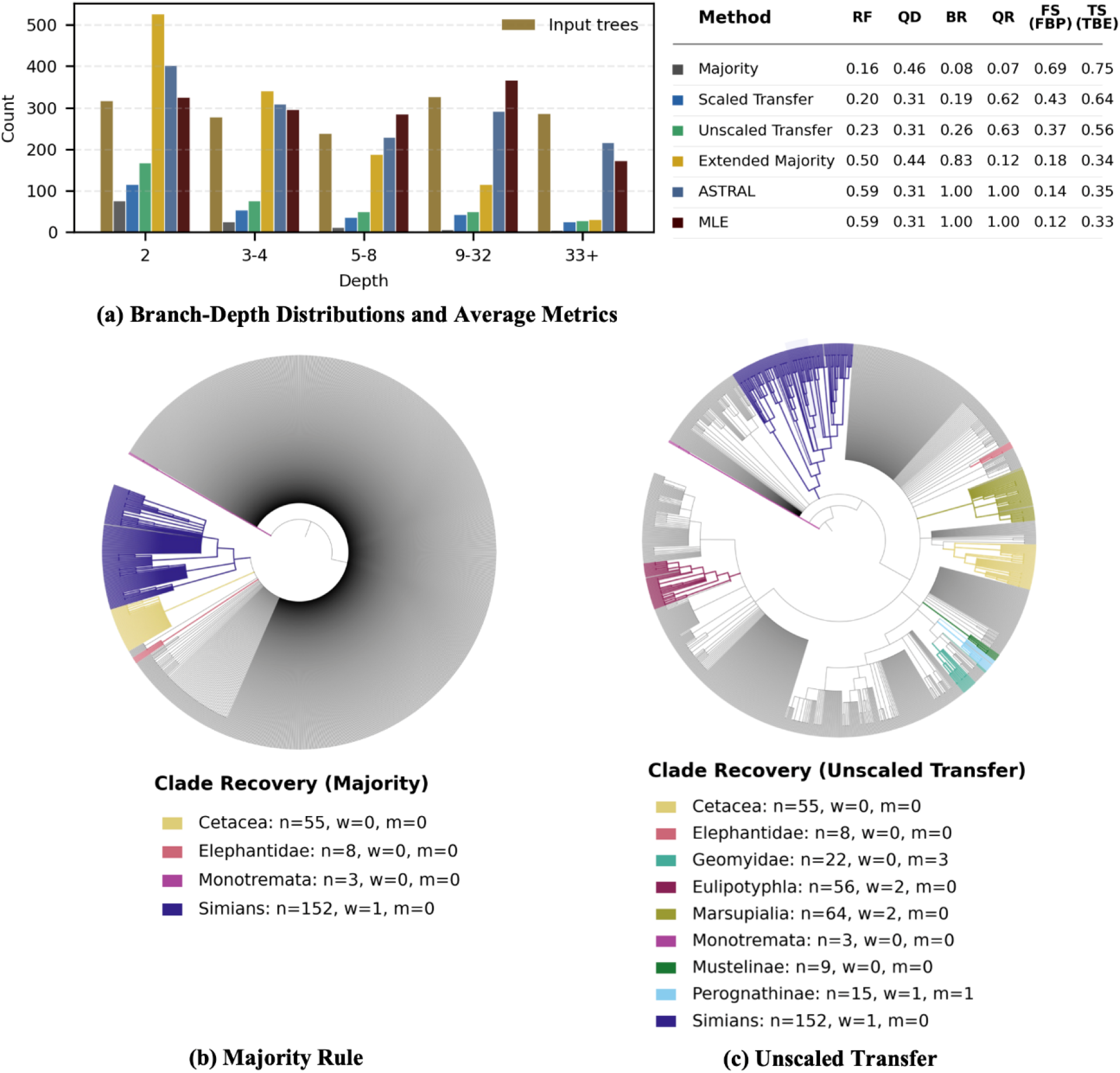
Full mammal dataset. **(a)** Distribution of branch depths (left) and summary statistics (right). **(b)** Majority-rule and **(c)** unscaled-transfer consensus trees computed from the full dataset (1,449 taxa). Highlighted clades correspond to major mammalian groups reported in Lemoine et al. (2018). A clade is considered approximately recovered if its closest matching clade has transfer distance error ≤15% (n: clade size; w: wrong taxa; m: missing taxa); when the error is zero (w = m = 0), the clade is fully recovered. The scaled-transfer consensus recovers the same nine reference clades under the 15% threshold and is not shown. The table summarizes metrics for the full-dataset analysis; RF: bipartition distance to the NCBI taxonomy; QD: normalized symmetric quartet difference, restricted to quartets resolved in the NCBI taxonomy; BR: branch resolution; QR: quartet resolution; FS (FBP): average frequency support (Felsenstein Bootstrap Proportion); TS (TBE): average transfer support (Transfer Bootstrap Expectation).

Results for the 100 subsampled datasets (164 taxa each), for which the NCBI taxonomy is fully resolved, are summarized in Figure 7. Consistent with the simulation experiments, the proposed consensus trees improve all three fine-grained dissimilarities relative to the majority-rule consensus, both with respect to the bootstrap input trees (results not shown) and the NCBI reference tree. The resolution gains are significant, with quartet resolution increasing by nearly threefold. Although they have many more branches than majority rule (more than twice as many with quartet-based consensus), they have relatively low bipartition loss (5%, 7%, and 14% for scaled-transfer, unscaled-transfer, and quartet-based consensus, respectively) while gaining ~20% in quartet-based topological accuracy with respect to the NCBI tree. As expected, this gain in resolution is also accompanied by lower average branch support than for majority rule (FBP: 0.47–0.59 versus 0.74; TBE: 0.63–0.74 versus 0.80). However, the branch support values are higher than those for EM, ASTRAL-IV, and MLE (FBP: 0.32– 0.37; TBE: 0.55–0.58). In terms of branch-depth distribution profiles (Fig. 7a), quartet-based approaches (our consensus and ASTRAL-IV) and EM tend to infer many shallow branches. Transfer-based consensuses do not exhibit any specific bias, and the majority rule infers very few deep branches (as in all of our analyses). As in the simulation experiments, the best-scoring highly resolved trees are either given by ASTRAL-IV or EM rather than MLE. This result again suggests the usefulness of the consensus of bootstrap trees over the original tree estimate.

**Figure 7.**
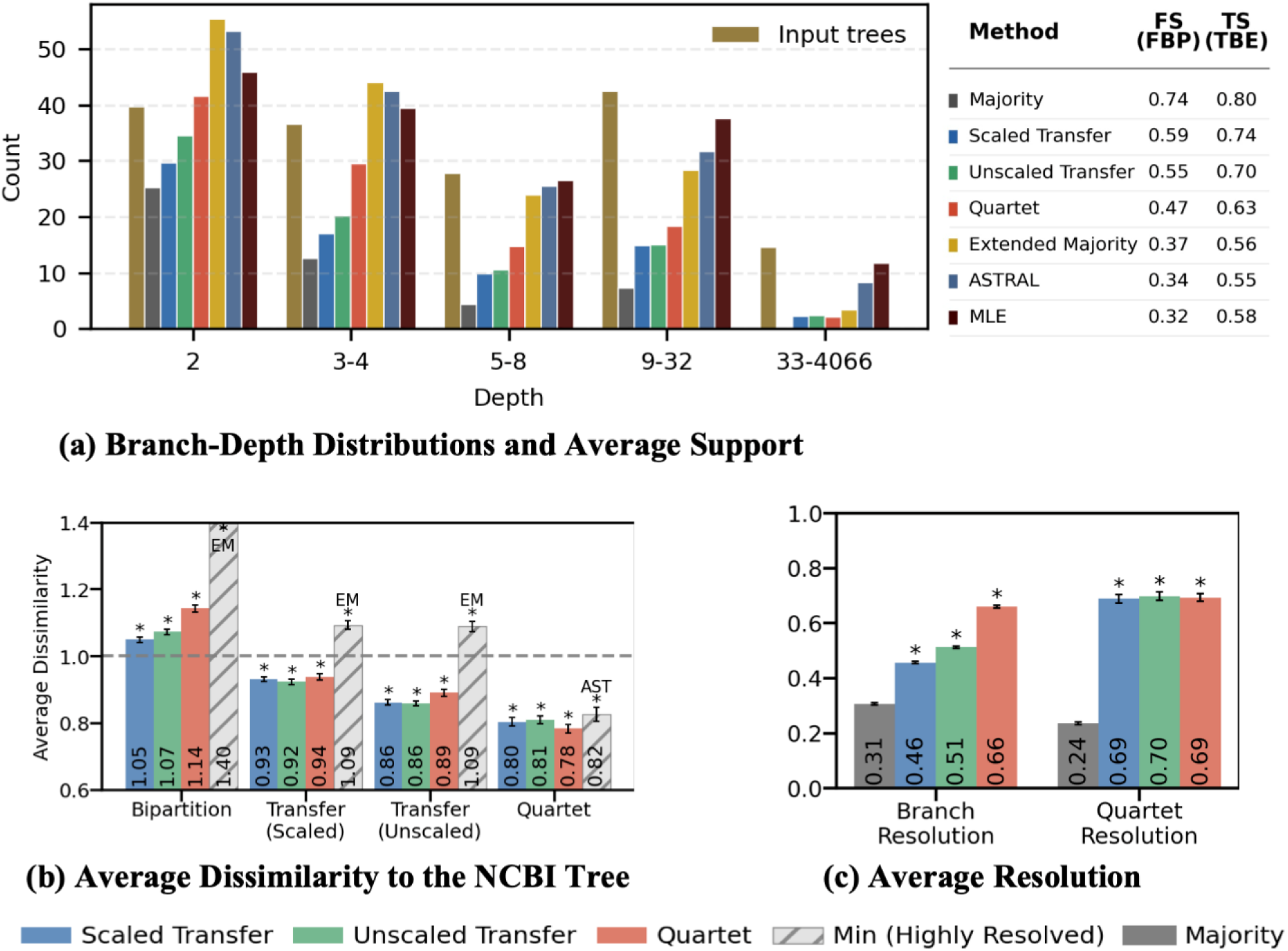
Subsampled mammal dataset. **(a)** Distribution of branch depths (left) and average support values (right). FS (FBP): average frequency support (Felsenstein bootstrap proportion); TS (TBE): average transfer support (transfer bootstrap expectation)). **(b)** average dissimilarity to the NCBI tree (Majority: dashed horizontal line, Y=1.0); **(c)** average branch and quartet resolution. The dashed gray bar in **(b)** indicates the minimum loss among highly resolved approaches (EM, AS: ASTRAL-IV, MLE). Error bars show 95% confidence intervals. Asterisks denote paired t-test significance relative to the majority-rule consensus (p < 0.01).

#### Analysis of HIV Data

The HIV dataset is very large. It comprises 1,000 bootstrap trees with over 9,000 leaves, taken from Lemoine et al. (2018). In this section, we aim to assess the applicability of our consensus methods, as well as their properties. We will also check whether these consensus methods can recover the deep structure corresponding to the nine HIV-1M subtypes. The quartet-based consensus method was not considered here, as it requires excessive computing time. In fact, even with the fast options, ASTRAL-IV itself required more than one week on a large cluster (see Supplementary Material for details). The methods studied are the majority-rule consensus, EM, our two transfer-based methods, ASTRAL-IV and MLE. Figure 8 summarizes the results.

**Figure 8.**
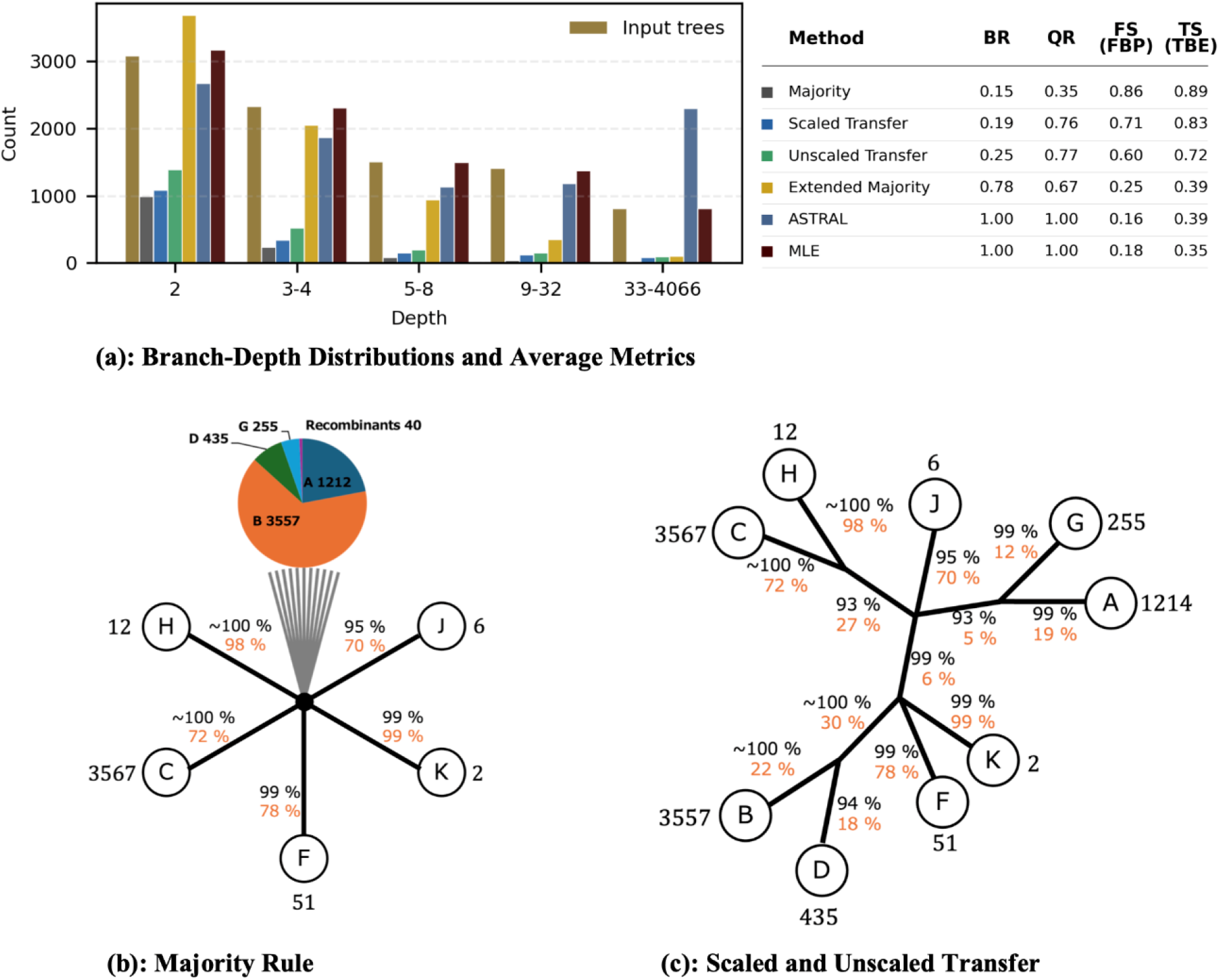
Comparison of consensus trees on the HIV dataset. **(a)** Distribution of branch depths (left) and summary statistics (right) for each consensus method; BR: branch resolution; QR: quartet resolution; FS (FBP): average frequency support (Felsenstein bootstrap proportion); TS (TBE): average transfer support (transfer bootstrap expectation). **(b, c**) Deep-branching structure of the majority-rule consensus tree **(b)** and the consensus trees obtained by minimizing the scaled- and unscaled-transfer dissimilarities **(c)**. In **(b, c)** the recombinants are not shown, and the displayed clades have a perfect match with the corresponding subtype, except for the recombinants. The numbers next to the clades indicate the number of samples of the corresponding subtype in the dataset. Values along branches report the FBP (orange) and TBE (black) supports of the associated split, computed from the 1,000 input bootstrap trees. For some branches, these two supports differ radically, with low FBP support but TBE support close to 100%. This corresponds to a few rogue or recombinant taxa with an unstable position relative to the given clade. For example, the B clade in **(c)** perfectly recovers the B subtype, but contains two recombinants which behave as rogue taxa and explain the low FBP support, while TBE is close to 100%. In (b), the central node has degree 4,109 in the original majority-rule tree; the gray fan represents the numerous incident branches omitted from the simplified drawing, and the pie chart summarizes the subtype composition within the highlighted clade, including 40 recombinants. Within this clade, the remaining subtypes (A, B, D, and G) are not even approximately recovered (see text and Supplementary Material for details).

The deep structure of the majority-rule consensus tree is star-like (Fig. 8b). Its average supports are high (FBP: 0.86, TBE: 0.89), but it contains only a small number of branches (BR: 0.15, QR: 0.35). It fails to recover several HIV-1M subtype clades, namely subtypes A, B, D, and G (*n* = 1,214, 3,557, 435, and 255, respectively). These four subtypes are not even approximately recovered when accepting a high fraction of errors (50% or more). Excluding C (*n* = 3,567), the recovered subtypes F, H, J, and K (which are perfectly matched, except for the recombinants) contain a small number of sequences/leaves (*n* = 51, 12, 6, and 2, respectively).

On the opposite end, ASTRAL-IV and MLE are fully resolved, while EM shows relatively high branch resolution, but medium quartet resolution (BR: 0.78; QR: 0.67). These three trees fully recover the nine subtypes (except for the recombinants) thanks to their high branch resolution. However, they are unreliable overall and have low branch support (TBE: 0.39 for EM/ASTRAL and 0.35 for MLE; FBP is substantially lower), as expected with such a low-signal dataset. In fact, these trees are radically different in terms of shape. The branch-depth distribution profiles (Fig. 8a) show that EM infers many cherries (2-leaf subtrees) and few deep branches, while ASTRAL-IV has a high number of deep branches, which typically corresponds to an unbalanced tree shape. MLE falls in between in terms of tree shape.

The two proposed transfer-based consensus trees offer a reasonable compromise between these extremes. They are conservative in branch resolution (BR: 0.19 and 0.25 for scaled- and unscaled-transfer, respectively) yet obtain substantially higher quartet resolution (QR: 0.76 and 0.77, respectively) than EM (QR: 0.67). This suggests that they prioritize well-supported deep structures over introducing numerous weakly supported, shallow clades (e.g., cherries). This is reflected in their higher average support relative to EM (TBE = 0.83 and 0.72 versus 0.39, with similar gaps regarding FBP). Importantly, the two proposed consensus trees fully recover the nine subtypes (except for the recombinants; Fig. 8c). Additionally, the deep branching of the subtypes is consistent with the literature (Hemelaar 2012), with TBE support approaching 100%. However, two of the deep nodes remain unresolved with a degree of 4.

Overall, Figure 8 highlights distinct failure modes on this dataset: majority rule can be so unresolved that it omits essential subtype-level structure, whereas highly resolved methods may introduce weakly supported or unstable deep relationships. In contrast, the proposed transfer-based consensuses retain comparatively strong support while recovering the deep subtype-defining structure, providing a more stable compromise for large trees.

In summary, applying our transfer-based consensus approach to this large, low-signal dataset demonstrates that major improvements can be achieved compared to the majority-rule consensus tree. Furthermore, computing the transfer-based consensuses of the 1,000 bootstrap trees by pruning the MLE tree, is achieved in a relatively short time of about 20 minutes on our laptop (MacBook Pro, Apple M2 Pro, 32 GB memory).

## Discussion and Conclusion

While the majority-rule consensus tree has been widely used to summarize multiple phylogenetic trees and has theoretical optimality as a median tree with respect to the bipartition (or Robinson-Foulds, RF) distance, it has one well-known drawback: it tends to produce highly unresolved trees with large datasets, making it difficult to gain biological insights. In this paper, we have proposed novel consensus tree methods to overcome this limitation. Each of our methods computes a median tree with respect to a fine-grained tree dissimilarity measure instead of the bipartition distance. Our focus was on three measures: scaled-transfer dissimilarity, unscaled-transfer dissimilarity, and quartet distance. We also provided fast algorithms for obtaining approximate median trees with respect to transfer-based dissimilarities.

Our simulation study confirmed improved tree resolution and loss reduction in both Bayesian and bootstrap settings, with particularly notable improvements in challenging scenarios with low phylogenetic signal. In the Bayesian setting, we observe a 6-8% increase in branch resolution and a 15-16% increase in quartet resolution. In the bootstrap setting, branch resolution improved from 45% to 55-65%, while quartet resolution increased dramatically from 41% to about 80%, corresponding to the recovery of deep branches in consensus trees. The Bayesian analysis results showed that our approaches effectively summarize the posterior distribution while providing reliable estimates of the true tree topology, as measured using our fine-grained dissimilarity measures. The reduction in loss compared to the input trees and the true tree is even higher in bootstrap analyses.

In addition to the main comparison with the majority rule, our simulation study showed that by comparing the performance of methods that produce highly resolved trees, EM and ASTRAL outperform conventional approaches such as MAP and MCC. This suggests that, in Bayesian settings, combining information from multiple posterior trees yields better summaries and tree estimates than methods that select a single posterior sample tree. The analysis of the Bayesian benchmark recently proposed by Berling et al. (2025) further reinforces this conclusion, demonstrating the superiority of the CCD-MAP approach (or ASTRAL or EM) over MCC. Similarly, in the bootstrap case, we found that EM and ASTRAL tended to outperform the MLE tree inferred from the original (non-resampled) multiple sequence alignment, in a manner similar to the bootstrap aggregating (bagging) approach used in machine learning (Breiman 1996). Overall, these results suggest that refined consensus methods, such as ours, should be considered for more than just summarizing sets of posterior or bootstrap trees. They should also be used to obtain more reliable tree estimates by leveraging the information provided by the variability of bootstrap or Bayesian posterior trees. It should be noted, however, that our consensus approaches are not intended to combine gene trees to obtain a species tree, as ASTRAL, for example.

We further validated our approaches by analyzing large real-world datasets, demonstrating both computational efficiency and biological relevance. For the West-African EBOV dataset, our consensus trees substantially improved branch and quartet resolution by ~30% and ~50%, respectively, compared to majority rule, while maintaining high average support values (FS: ~75%, TS: ~90%). Geographic analysis shows that our consensuses more accurately recover the structure associated with the regional spread of the virus, at a level similar to that of nearly or fully resolved approaches, while avoiding over-resolution and weak support values (FS: ~50% and TS: ~70%, in average). For the mammal dataset, our methods achieved better resolution than the majority rule while maintaining high accuracy. For example, analysis of the full mammal dataset (Fig. 6) showed that the majority-rule consensus tree is mostly unresolved (branch resolution: 8%) with a high quartet distance to the NCBI tree (46%). In contrast, the unscaled-transfer consensus tree has over three times as many branches (branch resolution: 26%) and significantly reduces the quartet-based topological error (31%), while the bipartition distance to the NCBI tree shows a moderate increase (23% vs 16% for majority rule). For the HIV-1M pol sequence dataset (9,147 taxa), where the true phylogeny is unknown, our consensus trees based on scaled- and unscaled-transfer dissimilarities showed higher resolution than the majority-rule consensus tree, particularly in terms of quartet resolution. Although not fully resolved, as expected given the number of taxa and low phylogenetic signal, our consensus trees successfully recovered all nine HIV-1M subtypes, while the majority-rule consensus tree failed to identify four subtypes. These results with large real-world datasets demonstrate that our fine-grained consensus methods can provide more meaningful biological insights than traditional majority-rule consensus trees.

In all of our experiments, but one expected case, the proposed consensus trees retained almost all (~98–100%) of the majority-rule bipartitions while adding further branches (see Supplementary Table S10 for details). This indicates that the increased resolution primarily extends the majority-rule tree. As expected from the inclusion of more branches, the additional resolution is accompanied by lower average support than for majority rule. However, the proposed consensus trees generally had high average transfer support, and the standard and transfer branch supports were substantially higher than those of the highly resolved alternatives. Furthermore, branch-depth analyses revealed no consistent propensity for the proposed methods to add branches that were disproportionately shallow or deep.

The Bayesian benchmark provides partial evidence regarding the stability of consensus estimation with finite samples. For both Coal320 and Yule400, disagreement between consensus trees obtained from two independent Markov chain Monte Carlo (MCMC) runs decreased as the number of posterior trees increased. Our consensus trees showed small between-run differences at a sample size of 1,000. However, this does not imply that 1,000 trees are universally sufficient; the required sample size may depend on factors such as the number of taxa, the phylogenetic signal and the variability of the input-tree distribution, as well as the mixing of the Markov chain Monte Carlo (MCMC) and the sampling dependence in Bayesian analyses. Moreover, the use of a single tree to summarize a posterior tree distribution may be limited, and several authors suggested using multiple trees with complex data (Bonnard et al., 2006; Silva and Wilkinson, 2021). Similarly, when using a consensus tree approach to summarize very large trees in a bootstrap setting, as with HIV, limitations arise (e.g., regarding branch support and local topological configuration), particularly when the phylogenetic signal is weak and the input trees differ substantially.

A comparative analysis of our three proposed consensus methods provides practical guidance for their application. Although they all performed comparably well in most experiments, computational constraints prevented us from approximating the quartet-distance-based median tree in the analysis of the full-mammal and HIV datasets. By contrast, the computational cost of the transfer-based approaches, particularly with pruning (Strategy 1), is negligible relative to Bayesian posterior sampling or bootstrap tree inference. Using a single thread on a laptop, the scaled- and unscaled-transfer pruning methods required only about 30 seconds on the Yule400 benchmark (Supplementary Table S11), while the HIV analysis with 9,147 taxa and 1,000 bootstrap trees required about 20 minutes. In addition to computational considerations, there are several important practical differences to consider when choosing the most appropriate method: (i) Each approach optimizes fundamentally different dissimilarity measures; the scaled-transfer dissimilarity most closely resembles the bipartition distance, assigning equal weights to all branches while replacing the exact match indicator with a gradual similarity measure in the [0,1] range, whereas weighted approaches, particularly the quartet-based method, place greater emphasis on deep branches. (ii) As a result, weighted approaches (quartet and unscaled-transfer) typically achieve higher resolution gains, especially in terms of quartet resolution, where deep branches significantly affect the measurements. (iii) While weighted approaches achieve higher branch resolution, they typically suffer from greater bipartition-distance loss when measured against the input trees. Consequently, the average level of support tends to be lower for the weighted approaches than for the unweighted ones. For example, in the subsampled mammal dataset, the mean transfer support across internal branches, averaged over the 100 replicates, was 80% and 74% for the majority rule and the scaled-transfer consensuses (unweighted approaches), compared to 70% and 63% for the unscaled-transfer and quartet methods (weighted approaches). On the other hand, our three consensus approaches infer a large number of deep branches, considerably improving the tree resolution by a factor of ~1.5 to ~2.0 for the branches and ~3.0 for the quartets (Fig. 7). If one is interested in the deep part of the tree, our consensuses provide essential information.

Our results suggest several important directions for future research: (i) Both theoretical and practical properties of these consensus trees should be further explored when combined with different tree inference methods, substitution models, data types, etc. (ii) A key issue is robustness to model misspecification when inferring the input trees, with the goal that the consensus tree is more robust than the individual input trees. Further results are needed along this line. (iii) The finite-sample stability of consensus trees should be investigated more systematically across datasets, sample sizes, and levels of phylogenetic signal. (iv) Although we have implemented efficient pruning algorithms for the two versions of the transfer-based consensus, such an algorithm is still needed for the quartet-based consensus. Exploring variants of transfer-based consensus would also be worthwhile, particularly with the goal of maximizing the transfer support of the consensus tree branches. Finally, algorithmic developments should be conducted to perform a more thorough search than the simple greedy tree-pruning approach and possibly obtain exact solutions.

## Acknowledgments

We thank Frédéric Lemoine and Paul Zaharias for their help with the mammal and HIV datasets. We also thank Krister M. Swenson for providing code for rapid computation of transfer bootstrap expectation. Finally, we thank the editors and anonymous referees for their comments that helped us improve the manuscript. Computations were partially performed on the NIG supercomputer at ROIS National Institute of Genetics and the UTokyo Azure platform (Nakamura et al. 2025).

## Data Availability

All the datasets used in this study can be accessed through Dryad (temporary Reviewer URL for review; DOI for accepted manuscript) and GitHub https://github.com/yukiregista/PhyloCRISP.

## Supplementary Material

## A Maximum of the unscaled transfer dissimilarity

In this section, we prove the following proposition regarding the maximum possible value of the unscaled transfer dissimilarity between two trees with *n* taxa. This result is relevant for understanding the range and extremal behavior of the measure. Note that a *caterpillar tree* is a tree in which there exist two leaves such that all nodes are either on the path between them or adjacent to a node on that path.

### Proposition 1.

*The maximum unscaled transfer dissimilarity between two trees with n taxa is n*^2^*/*2 − 2*n* + 2 *when n is even and n*^2^*/*2 − 2*n* + 3*/*2 *when n is odd*.

*Proof*. First, we consider the maximum sum of depths of internal bipartitions in a tree with *n* taxa. Recall that the depth of a bipartition in this paper refers to the cardinality of the smaller part. [6] derived that for any strictly increasing function *f* : {2, …, ?*n/*2?} → R_*>*0_, the following type of quantity is (only) maximized by caterpillar trees:

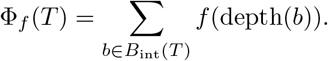

In particular, choosing *f* (*x*) = *x* corresponds to summing the depths of all internal bipartitions, which is directly related to the transfer distance calculation. In a caterpillar tree on *n* taxa, the internal bipartitions have depths min(*i, n* − *i*) for *i* = 2, …, *n* − 2. Therefore, the maximum sum is achieved by caterpillar trees and is given by

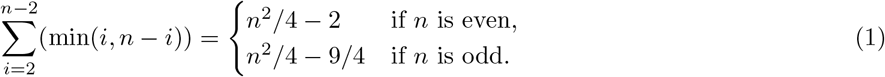

Recall that the unscaled transfer dissimilarity between two trees *T*_1_ and *T*_2_ with *n* leaves is defined as follows. For each internal bipartition in one tree, we compute the minimum transfer distance to any bipartition in the other tree, and sum these values over all internal bipartitions in both trees:

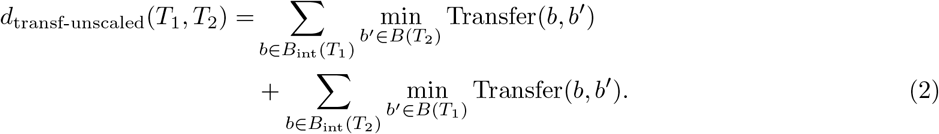

Note that each summand cannot be larger than depth(*b*) − 1 since the transfer distance between *b* and some appropriate external branches is equal to that amount. Combining (1) and (2), we can bound the unscaled transfer dissimilarity as follows:

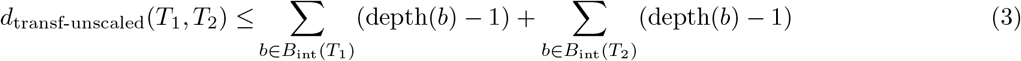

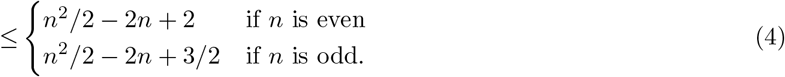

It remains to show that the bound in (4) is attainable. Let

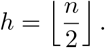

We construct two caterpillar trees with taxon orderings *σ*_1_, *σ*_2_ and binary functions *f*_1_, *f*_2_ such that

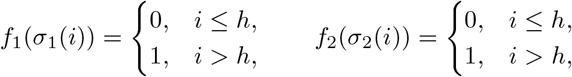

while

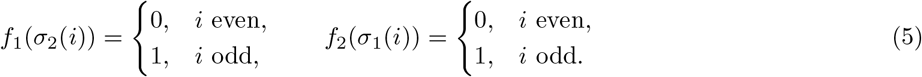

Thus each function indicates the two halves of one caterpillar and alternates along the other caterpillar.

Such a pair is easily constructed. Take *σ*_1_ to be the identity, *f*_1_(*x*) = 0 for *x* ≤ *h* and 1 otherwise, and let *f*_2_(*x*) = 0 for even *x* and 1 for odd *x*. Construct *σ*_2_ by placing all even taxa in its first *h* positions and all odd taxa thereafter, with taxa greater than *h* in odd positions and taxa at most *h* in even positions. The required numbers in these four classes agree, so this defines a permutation satisfying (5).

Let *A* and *B* be the smaller parts of bipartitions of the two caterpillars, with |*A*| = *k*, |*B*| = *m*, and assume *k* ≤ *m*. Since *f*_2_ is constant on *B* and alternates along *A*, if *β* is the value of *f*_2_ on *B*, then

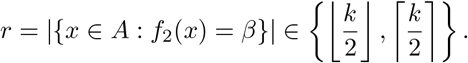

Writing *c* = |*A* ∩ *B*|, we have *c* ≤ *r*, and therefore

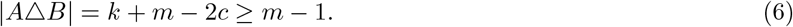

Let *s*_*β*_ = *x* : *f*_2_(*x*) = *β*. Since *B* contains *m* elements of this class, at most *s*_*β*_ − *m* of its elements lie outside *B*. Hence

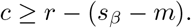

Therefore,

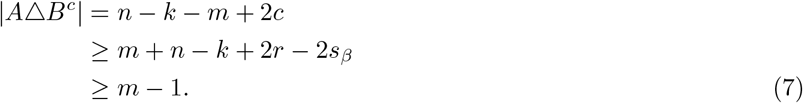

The last inequality follows directly from the alternating construction: if *n* is even, *s*_*β*_ = *n/*2 and *r* ≥ ?*k/*2?; if *n* is odd, the two class sizes are ?*n/*2? and ⌈*n/*2⌉, with the corresponding counts in *A* equal to ?*k/*2? and ⌈*k/*2⌉.

Since

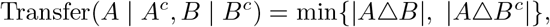

(6)–(7) imply

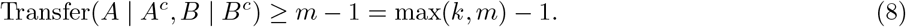

The case *m < k* follows by symmetry.

Consequently, for every internal bipartition *b* of either caterpillar, every bipartition of the other tree is at transfer distance at least depth(*b*) − 1. This bound is attained by an external bipartition corresponding to any taxon in the smaller part of *b*. Hence

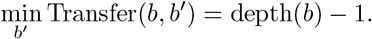

Since both trees are caterpillar trees, equality is attained in (3), and therefore the upper bound in (4) is tight.

## B Details on the Pruning Algorithm

### B.1 Pruning with respect to transfer-distance-based dissimilarity: algorithm overview

We present a pruning algorithm for efficiently computing an approximate median tree with respect to the scaled transfer dissimilarity. The same framework applies, with minor modifications, to the unscaled transfer dissimilarity. The algorithm iteratively prunes branches from an initial tree to minimize the overall dissimilarity to a set of input trees.

#### Inputs and Outputs

- **Input:** An initial binary tree *T*_0_, a set of input trees *T* = {*T*_1_, …, *T*_*N*_}, and a hyperparameter *K* ≥ 2 (the number of candidate matches retained for each input-tree bipartition).
- **Output:** The set of remaining bipartitions *E* corresponding to the pruned consensus tree obtained from *T*_0_.

#### Algorithm Structure

The pruning algorithm consists of three main steps:

1. **Computation of transfer distances and support values:** For each bipartition in the initial tree, compute its transfer support with respect to the input trees, and vice versa.
2. **Initialization:** Set up arrays to track the contributions of each bipartition to the false positive and false negative components of the loss function.
3. **Iterative pruning:** Repeatedly prune the branch that yields the greatest reduction in loss, updating relevant data structures at each step.

#### Objective Function

Recall that the objective function to be minimized is decomposed into false positive and false negative components:

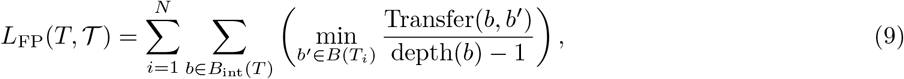

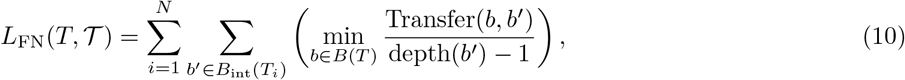

where *B*(*T*) denotes the set of all bipartitions in tree *T*, and *B*_int_(*T*) denotes the set of internal bipartitions. The false positive term penalizes branches in *T* not well supported by the input trees, while the false negative term penalizes bipartitions in the input trees that do not have a similar (i.e., low transfer distance) counterpart in *T*.

#### Step 1: Computation of Transfer Support and Best Matches

First, we compute the transfer support values for each bipartition in the initial tree *T*_0_ with respect to the input trees *T*. These values determine the contribution of each branch *b* to the false positive component *L*_FP_, which remains constant throughout pruning. However, the sum of these decreases along the pruning process, where branches in *T* are removed.

Next, for each input tree, we compute the support of each of its bipartitions with respect to *T*_0_. This step allows us to evaluate the false negative loss *L*_FN_ for *T*_0_. Unlike the false positive component, the contribution of each bipartition to the false negative loss can change as pruning progresses, since the best-matching bipartition in *T* (starting from *T*_0_) may change after each pruning step.

To efficiently track these changes, we precompute the *K* closest bipartitions in *T*_0_ for each bipartition in the input trees. This precomputation enables rapid calculation of the change in transfer distance when the best match is replaced by the second-best match, which directly determines the change in the false negative loss. As long as the updated best matches remain within the initial top *K* candidates, we can update the transfer distances and loss values efficiently without recomputing all pairwise distances.

#### Step 2: Initialization

We initialize the following arrays:

- FP: Stores the false positive loss for each bipartition.
- *δ*_FN_: Stores the potential increase in false negative loss if a bipartition is pruned.
- *L* = FP − *δ*_FN_: Represents the net benefit of pruning each branch.

We also initialize data structures to track the current best and second-best matches for each bipartition, as well as their associated ranks.

#### Step 3: Iterative Pruning

The iterative pruning process proceeds as follows:

- While there exists a branch with *L >* 0, select the branch *b* with the largest *L* value.
- Prune branch *b* from the current tree *T*.
- Update all relevant data structures, including the best and second-best matches for affected bipartitions.
- If the set of precomputed *K* matches is exhausted for any bipartition, recompute transfer distances to all remaining branches as needed.

### B.2 Pruning with respect to transfer-distance-based dissimilarity: details

Algorithm 1 summarizes the pruning procedure. We now detail each of its steps.

#### B.2.1 Computation of transfer support and hash values

The function TbeSupport(*T*_0_, *T*) computes the transfer support for each bipartition in *T*_0_ with respect to the input trees *T*, using the algorithm of [15] in *O*(*nN* (log *n*)^3^) time. In addition to support values, we compute hash values for efficient bipartition lookup and track the frequency of each bipartition across the input trees.

For hashing, we use the method of [1], which employs two universal hash functions. To define the hash, we first represent each bipartition as a bitstring of length *n*, where the *i*-th bit indicates the side of taxon *i*; a unique bitstring can be assigned for each bipartition, for example, by consistently assigning a fixed value to the bit corresponding to a chosen taxon. Given the bitstring representation of a bipartition, *b*_1_, …, *b*_*n*_, two prime numbers *M*_1_ and *M*_2_, and nonnegative integer coefficients 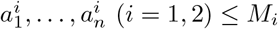, the hash values are:

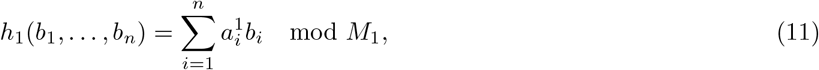

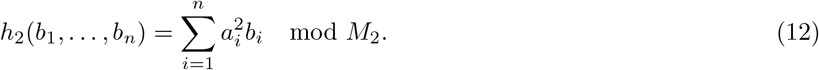

We use an array of size *M*_1_ indexed by *h*_1_, with each entry a linked list of *h*_2_ values. The combined hash (*h*_1_, *h*_2_) has collision probability 1*/*(*M*_1_*M*_2_). The memory requirement is *O*(*M*_1_ + *m*), and *M*_2_ can be chosen large (e.g., 2^31^ − 1). Hashing all bipartitions in an input tree takes *O*(*n*) time. We also store the frequency and tree ID for each bipartition.

Currently, we do not check for hash collisions, so a collision between two distinct bipartitions occurs with probability approximately 1*/*(*M*_1_*M*_2_) under the hashing scheme. We choose *M*_1_ = *O*(*nN*), yielding time and memory complexity *O*(*nN* (log *n*)^3^) and *O*(*nN*), respectively. As a safe alternative, we could store bitstrings and check for collisions, at the cost of *O*(*n*^2^*N*) time and memory.

The output of TbeSupport is:

- TS: Array of size *n* − 3, transfer support for each internal bipartition in *T*_0_.
- *H*_1_, *H*_2_: Arrays of size *m*, hash values for bipartitions.
- *I*: Array of size *m*, storing for each unique bipartition one representative input-tree ID in which it occurs.
- *C*: Array of size *m*, occurrence counts for each bipartition in *T*.

##### Algorithm 1

Greedy Pruning Algorithm (Scaled Transfer)

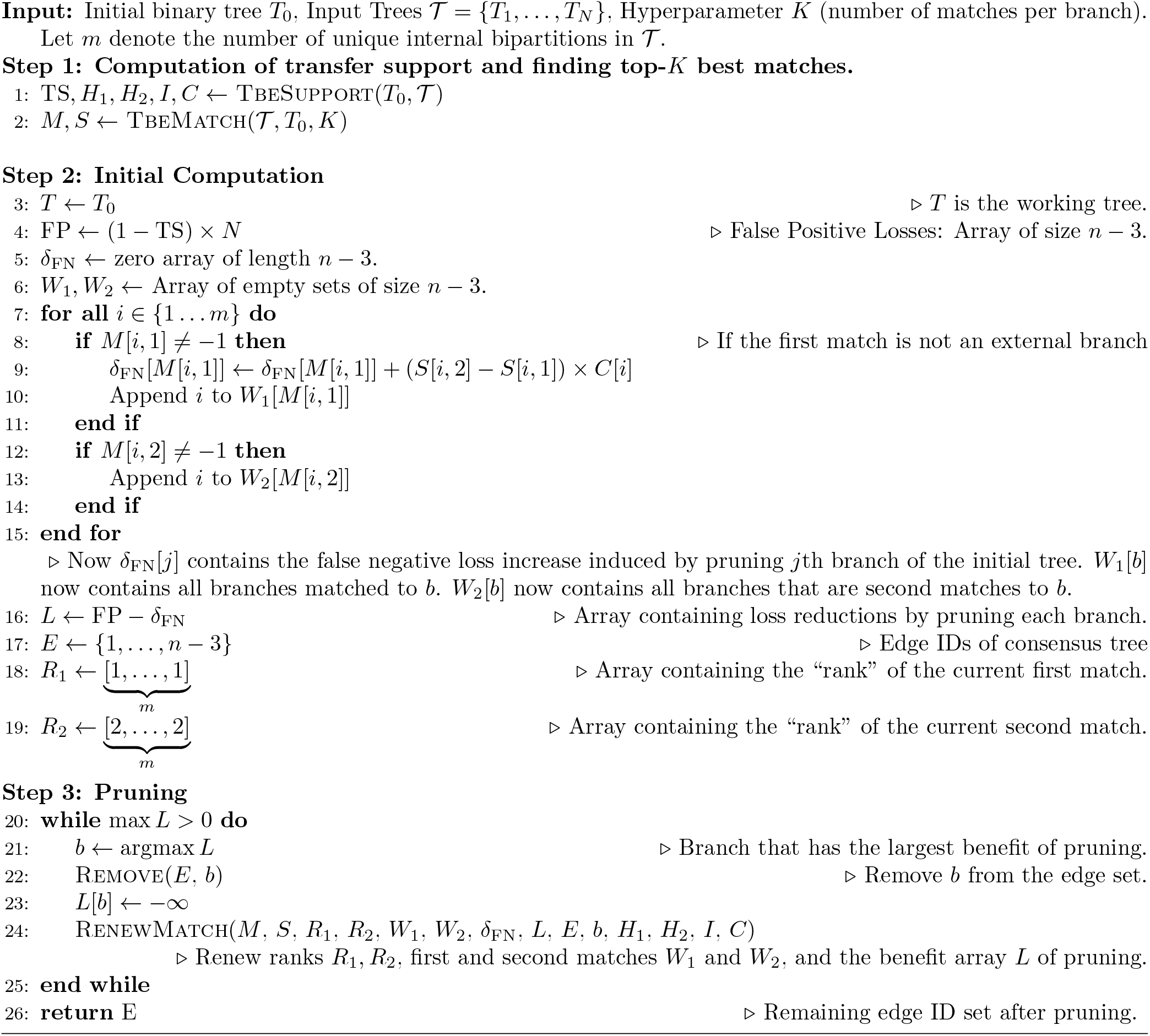

#### B.2.2 Computation of first *K* best matches

The function TbeMatch(T, *T*_0_, *K*) considers each input tree as the reference and, for each bipartition in the input trees, returns the top *K* matches from *T*_0_ in terms of transfer distance. This is computed efficiently by modifying the algorithm of [15] to track the top *K* matches, running in *O*(*nNK*(log *n*)^3^) time.

The output consists of:

*M* : Array of size *m* × *K*, where (*i, j*) stores the *j*-th best matching bipartition from *T*_0_ to the *i*-th bipartition from *T*.

*S*: Array of size *m* × *K*, storing the scaled transfer distance for each match.

Entries in *S* greater than or equal to 1 are set to 1, and the corresponding entries in *M* are set to −1, where −1 denotes a match to an external branch, since the minimum scaled transfer distance to an external branch is always 1.

#### B.2.3 Initial computation

In step 2 of Algorithm 1, we initialize the candidate tree *T* to *T*_0_ and compute the initial benefit of pruning each internal branch. The contribution of each internal branch *i* to the false positive loss is (1 − TS[*i*]) × *N*, stored in FP.

The increase in false negative loss induced by pruning branch *i* is denoted *δ*_FN_[*i*]. For a bipartition *b* in the input trees matched to *i* in *T*_0_, the scaled transfer distance is *S*[*b*, 1]; the second closest match is *S*[*b*, 2]. Pruning *i* increases the false negative loss for *b* by (*S*[*b*, 2] − *S*[*b*, 1]) × *C*[*b*]. Thus, 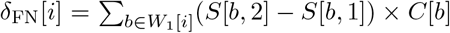, where *W*_1_[*i*] is the set of bipartitions in the input trees whose first match is *i*. We also maintain *W*_2_[*i*], the set whose second match is *i*. To compute the array *δ*_FN_, we initialize *δ*_FN_[*i*] = 0 for all *i*. Then, for each bipartition *b* in the input trees, we add (*S*[*b*, 2] − *S*[*b*, 1]) × *C*[*b*] to *δ*_FN_[*i*], where *i* is the first match of *b*, provided the first match is not an external branch.

We define *L* = FP − *δ*_FN_, representing the benefit of pruning each branch. Arrays *R*_1_ and *R*_2_, each of length *m*, are used to track the rank of the current first and second matches for each bipartition, respectively. Here, the rank indicates the position of the matching bipartition in the original sorted list of candidate matches from *T*_0_, with rank 1 corresponding to the closest match, rank 2 to the second closest, and so forth. At the initialization step, all elements of *R*_1_ are set to 1 and all elements of *R*_2_ are set to 2.

#### B.2.4 Pruning and updating matches

After completing the initialization, we iteratively prune the branch with the highest *L* value, as long as this value remains positive. Each pruning step requires careful updates to the data structures that track the matching between bipartitions in the input trees and those in the current tree. In particular, after pruning a branch, the contribution to the false negative loss, *δ*_FN_, and the *L* array must be updated, since some bipartitions may have had the pruned branch as their first or second match, which becomes unavailable. Consequently, the arrays *R*_1_, *R*_2_, *W*_1_, and *W*_2_ must also be updated to reflect these changes. This update process is performed by the RenewMatch procedure, which consists of the following steps, where *i* denotes the branch pruned at the current iteration:

1. **Update for bipartitions in** *W*_1_[*i*]: For each bipartition *b* in *W*_1_[*i*] (i.e., those whose current first match is branch *i*), promote their second match to become the new first match. This is achieved by appending *b* to *W*_1_[*M* [*b, R*_2_[*b*]]], removing *b* from *W*_2_[*M* [*b, R*_2_[*b*]]], and updating *R*_1_[*b*] to *R*_2_[*b*].
2. **Find the new second match:** After promoting the second match, a new second match for *b* must be found. Starting from rank *R*_2_[*b*] + 1, search for the next available bipartition in the sorted list of candidate matches from *T*_0_ that has not been pruned. Once this new match is found (at rank *R*_2_[*b*] + *l*), append *b* to *W*_2_[*M* [*b, R*_2_[*b*] + *l*]] and set *R*_2_[*b*] = *R*_2_[*b*] + *l*.
3. **Update the false negative loss:** After updating the first and second matches for each *b* ∈ *W*_1_[*i*], adjust *δ*_FN_ accordingly. Specifically, update *δ*_FN_[*M* [*b, R*_1_[*b*]]] by adding the difference in transfer distances between the new first and new second match, multiplied by the number of occurrences, i.e., (*S*[*b, R*_2_[*b*]] − *S*[*b, R*_1_[*b*]]) × *C*[*b*].
4. **Update for bipartitions in** *W*_2_[*i*]: Similarly, for each bipartition *b* in *W*_2_[*i*] (those whose current second match is branch *i*), search for the next available second match starting from rank *R*_2_[*b*] + 1. Once the new match is found (at rank *R*_2_[*b*] +*l*), append *b* to *W*_2_[*M* [*b, R*_2_[*b*] +*l*]] and update *R*_2_[*b*] to *R*_2_[*b*] +*l*. Adjust *δ*_FN_ for these bipartitions by updating *δ*_FN_[*M* [*b, R*_1_[*b*]]] with the difference in transfer distances between the previous and new second match, multiplied by the number of occurrences, i.e., (*S*[*b, R*_2_[*b*]] − *S*[*b, R*_2_[*b*] − *l*]) × *C*[*b*].

In some cases, it may be necessary to search beyond the *K*-th match, which requires computing additional matches beyond the originally precomputed top *K*. When this occurs, we recompute transfer distances between the relevant bipartition and all branches in *T*_0_ (or the current *T*) in *O*(*n*) time using the algorithm of [9], sort them in *O*(*n* log *n*) time, and store the results as an array. To retrieve the full representation of the bipartition, we use the hash values *H*_1_, *H*_2_, and the tree index *I*. This recomputation is performed at most once per bipartition.

If we ignore the time required by these additional computations and for updating matches beyond the *K*-th, the update process takes *O*(*K*) time per bipartition across all iterations, resulting in a total time complexity of *O*(*nNK*) for all *m* = *O*(*nN*) bipartitions. If only *O*(*NK*(log *n*)^2^) branches require recomputation, the total running time can be as fast as *O*(*nNK*(log *n*)^3^). In the worst case, where recomputation is performed for *O*(*nN*) bipartitions, the additional cost is *O*(*n*^2^*N* log *n*). The memory requirement is *O*(*nNK*) if we ignore the memory used for storing the results of recomputation; in the worst case, the total memory usage can be as large as *O*(*n*^2^*N*).

### B.3 Pruning with respect to quartet distance

To approximate the median tree under the quartet distance, we implement a greedy pruning algorithm that iteratively removes the branch whose deletion yields the largest decrease in symmetric quartet distance loss.

The *symmetric quartet distance* is defined as:

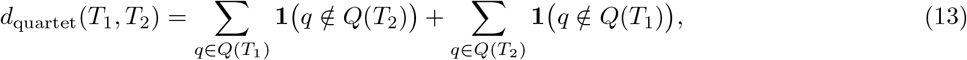

where *Q*(*T*) denotes the set of resolved quartet topologies in tree *T*. This symmetric form penalizes discordant resolved quartets more heavily than disagreements involving unresolved quartets, reflecting their greater phylogenetic significance.

We use tqDist [14] to compute the standard quartet distance and the number of unresolved quartets, combining these to evaluate the symmetric quartet distance loss at each pruning step.

To evaluate the loss reduction from pruning a branch *e*, we compare the quartet distance loss of the original tree *T* with that of the pruned tree *T*_−*e*_. A direct implementation recomputes this quantity for all branches at each step. However, this process can be greatly accelerated by observing that pruning a branch *e* only changes quartet sets uniquely supported by branches adjacent to *e*. Specifically, a quartet topology *AB* | *CD* is said to be *uniquely supported* by branch *e* in tree *T* if:

1. *A, B* lie on one side of the bipartition *b*_*e*_ induced by *e*,
2. *C, D* lie on the opposite side,
3. Within each side, *A* and *B* (resp. *C* and *D*) come from different subtrees incident to *e*.

Pruning *e* renders exactly these quartets unresolved, and the corresponding loss change can be computed accordingly. Moreover, when pruning *e* from *T*_*s*_ to obtain *T*_*s*+1_, any branch *f* not adjacent to *e* retains its surrounding subtree structure and bipartition. Hence, the set of quartets uniquely supported by *f* remains unchanged, and its loss-reduction score need not be updated. Only branches adjacent to *e* require recalculation, which significantly reduces the computational burden in practice.

## C Computational Methods and Software Configurations

### C.1 Consensus tree estimation

Across all simulation and empirical data experiments, consensus trees were constructed using the following software and parameter configurations:

#### Majority Rule Consensus

Implemented in Python using the DendroPy (version 5.0.8; https://jeetsukumaran.github.io/DendroPy) [11].

#### Extended Majority Rule Consensus

Performed using IQ-TREE2 (version 2.4.0; https://github.com/iqtree/iqtree2) [10]. The command used was:

~~~
iqtree2 -con <input_trees> -minsup 0
~~~

#### ASTRAL-IV

Executed using ASTER (version 1.22; https://github.com/chaoszhang/ASTER) [16] with the following command:

~~~
astral4 -i <input_trees> -o <output>
~~~

For the HIV-1M pol gene dataset, the options -r 1 -s 1 were appended to the command to alleviate computational constraints.

#### Maximum Clade Credibility (MCC)

Implemented in Python using DendroPy.

#### CCD0-MAP [4]

Performed using CCD package for BEAST2 (version 2.7.7; https://www.beast2.org) [5] with the following command:

~~~
treeannotator -topology CCD0 -burnin 0 <input_trees> <output>
~~~

#### HIPSTR [2]

For the EBOV analysis, HIPSTR was computed from the rooted posterior trees using TreeAnnotator in BEAST X [3] (version 10.5.0) with the following command:

treeannotator -type hipstr -heights mean -burninTrees 0 -threads 1 <input_trees> <output>

### C.2 Simulation experiments

We detail below the software versions and parameter settings used in the experiments.

#### C.2.1 Tree simulation

Birth–death trees were simulated using the Python package DendroPy with the following parameters:

~~~
birth rate=0.2, death rate=0.1, gsa ntax=1000, num extant tips=100, and is assign extinct taxa=False.
~~~

After sampling, a custom Python script (using DendroPy) was used to normalize tree height, perturb branch lengths by multiplying with log-normal random variables, and set a minimum branch length of 0.5*/*sequence length.

#### C.2.2 Sequence simulation (Seq-Gen)

Sequence evolution was simulated with Seq-Gen (version 1.3.5; https://github.com/rambaut/Seq-Gen) [12] using the HKY model. The command used was:

~~~
seq-gen -mHKY -n1 -l<length> -t2.0 -a0.5 -f0.15,0.35,0.35,0.15 -on -z<seed>
~~~

#### C.2.3 Maximum likelihood inference (RAxML-NG)

Maximum likelihood inference was performed using RAxML-NG (version 1.2.2; https://github.com/amkozlov/raxml-ng) [7] with the following command:

~~~
raxml-ng --msa <alignment> --model GTR+G --prefix MLE --seed <seed>
~~~

Bootstrap analysis used:

~~~
raxml-ng --bootstrap --blopt nr_safe --msa <alignment> \
--model GTR+G --bs-trees 1000 --seed <seed>
~~~

#### C.2.4 Bayesian inference (MrBayes v3.2.7a)

Bayesian inference was conducted using MrBayes (version 3.2.7a; https://github.com/NBISweden/MrBayes) [13] with the following configuration:

~~~
begin mrbayes;
set autoclose=yes nowarnings=yes seed=<seed> swapseed=<swapseed>; execute <alignment>;
lset nst=6 rates=gamma;
mcmcp ngen=1200000 burnin=201 relburnin=no
samplefreq=1000 nruns=4 nchains=4 temp=0.1 filename=<output>;
mcmc;
sump;
sumt;
end;
~~~

The four runs were used to assess convergence. Downstream analyses used the retained posterior trees from run 1 only.

### C.3 Real data experiments

#### C.3.1 Maximum likelihood inference for the subsampled mammal datasets (RAxML-NG)

Maximum likelihood inference with bootstrapping was performed using RAxML-NG (version 1.2.2; https://github.com/amkozlov/raxml-ng) with the following command:

~~~
raxml-ng --all --bs-trees 1000 --msa <alignment> --model mtMAM+G --seed <seed> --blopt nr_safe
~~~

#### C.3.2 Subtype matching criteria

In both the full mammal and HIV-1M pol gene dataset experiments, we evaluated how well the consensus trees recovered known reference clades (specifically, the annotated clades from [8] for the mammal data, and established viral subtypes for the HIV data). For a given consensus tree *T* and each reference clade bipartition *b*_ref_, we identified the bipartition *b*_match_ ∈ *B*(*T*) that minimizes the transfer distance to *b*_ref_. The clade recovery error *E* for that reference clade relative to the consensus tree is then defined as:

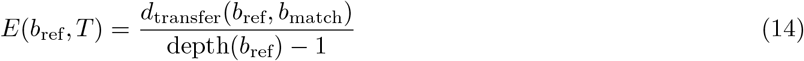

Moreover, for each best-matching clade, we also recorded the number of missing taxa from the reference clade and the number of “wrong” taxa included in the matched clade, where “wrong” taxa are taxa assigned to the best-matching clade but not belonging to the reference clade (see Figure 6 in the main text).

## D Detailed results of the experiments

### D.1 Performance of the majority-rule consensus tree

All the dissimilarity results given in the main text are normalized by the performance of the majority-rule consensus. In this subsection, we display the performance of the majority-rule consensus tree in Tables S1–S6. All dissimilarities are normalized in the following way: the bipartition distance, the scaled dissimilarity, and the unscaled dissimilarity are divided by 2(*n* − 3), where *n* is the number of taxa, and the quartet distance is divided by 2 ^*n*^. The unscaled dissimilarity can alternatively be normalized to the range [0, 1] using the result from Section A. Here, we normalize by 2(*n* − 3), motivated by the binary tree case: for two fully resolved trees, this yields the mean number of transfers required per branch. Although the majority-rule consensus tree is far from binary in this study, this normalization provides a scale that can be interpreted in relation to the fully resolved case.

**Table S1:**
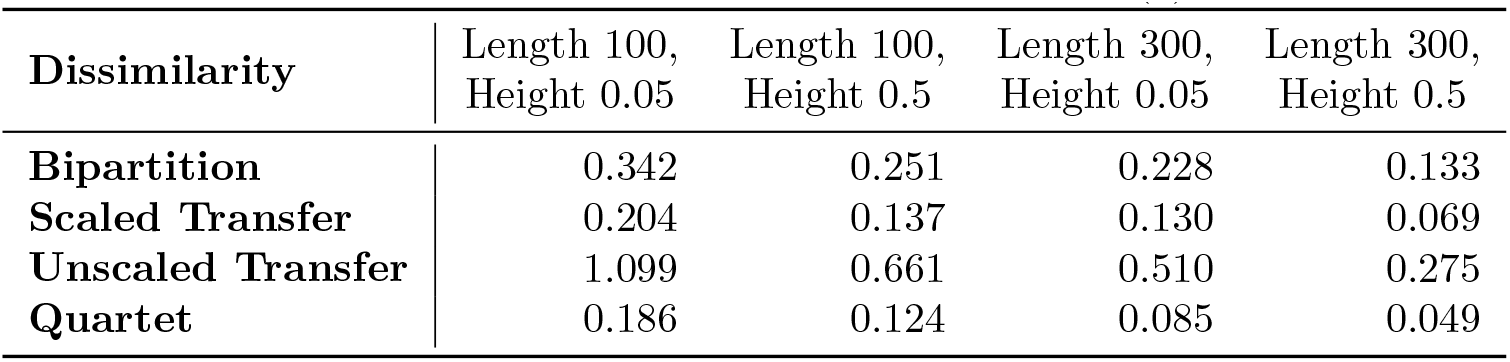
The normalized average dissimilarity between the majority-rule consensus tree and the **input** trees in the **Bayesian simulation experiments**, used as baseline values in Figure 1(a) in the main text.

**Table S2:**
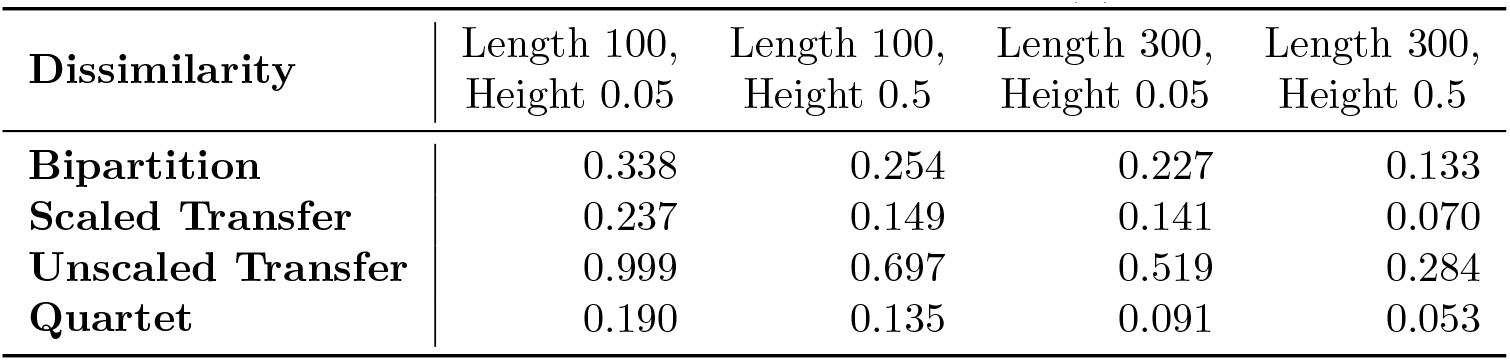
The normalized average dissimilarity between the majority-rule consensus tree and the **true** trees in the **Bayesian simulation experiments**, used as baseline values in Figure 1(b) in the main text.

**Table S3:** The normalized average dissimilarity between the majority-rule consensus tree and the input trees in the bootstrap simulation experiments, used as baseline values in Figure 3(a) in the main text.

| Dissimilarity | Length 100,<br>Height 0.05 | Length 100,<br>Height 0.5 | Length 300,<br>Height 0.05 | Length 300,<br>Height 0.5 |
| --- | --- | --- | --- | --- |
| **Bipartition** | 0.314 | 0.323 | 0.257 | 0.175 |
| **Scaled Transfer** | 0.190 | 0.185 | 0.135 | 0.088 |
| **Unscaled Transfer** | 1.729 | 1.299 | 0.771 | 0.423 |
| **Quartet** | 0.323 | 0.252 | 0.150 | 0.082 |

**Table S4:** The normalized average dissimilarity between the majority-rule consensus tree and the true trees in the bootstrap simulation experiments, used as baseline values in Figure 3(b) in the main text.

| Dissimilarity | Length 100,<br>Height 0.05 | Length 100,<br>Height 0.5 | Length 300,<br>Height 0.05 | Length 300,<br>Height 0.5 |
| --- | --- | --- | --- | --- |
| **Bipartition** | 0.377 | 0.268 | 0.245 | 0.141 |
| **Scaled Transfer** | 0.275 | 0.172 | 0.154 | 0.075 |
| **Unscaled Transfer** | 1.404 | 1.018 | 0.643 | 0.326 |
| **Quartet** | 0.300 | 0.226 | 0.124 | 0.064 |

**Table S5:** The normalized average dissimilarity between the majority-rule consensus tree and the true trees in the Bayesian benchmark experiment, used as baseline values in Figure 4(a, b) in the main text.

| **Dissimilarity** | Coal320 | Yule400 |
| --- | --- | --- |
| **Bipartition** | 0.247 | 0.141 |
| **Scaled Transfer** | 0.122 | 0.075 |
| **Unscaled Transfer** | 0.591 | 0.415 |
| **Quartet** | 0.034 | 0.055 |

**Table S6:** The normalized average dissimilarity between the majority-rule consensus tree and the NCBI tree in the bootstrap experiments with the subsampled mammal dataset, used as baseline values in Figure 7 in the main text.

| **Dissimilarity** | to NCBI |
| --- | --- |
| Bipartition | 0.399 |
| Scaled Transfer | 0.351 |
| Unscaled Transfer | 1.749 |
| Quartet | 0.383 |

### D.2 Comparison of the highly resolved consensus approaches in the simulation experiments

In this subsection, we show the performance of all highly resolved approaches (the extended majority (EM), ASTRAL-IV, MAP, MCC for the Bayesian case, and EM, ASTRAL-IV, MLE for the bootstrap case) in detail. Figure S1 and Figure S2 compare the performance of these highly resolved approaches in the Bayesian simulation experiment and the bootstrap simulation experiment, respectively.

For the Bayesian case, MAP is the worst approach in all settings. In addition, MCC is also worse than ASTRAL-IV and EM in all cases. This suggests that commonly used MAP and MCC are not suitable for the consensus of posterior draws in terms of tree topology metrics. The best approach is either ASTRAL-IV or EM, for both as a summary of posterior draws (average dissimilarity against input trees) and as a tree estimate of the true phylogeny. Likewise, in the bootstrap case, MLE is mostly worse than EM and ASTRAL-IV. These results suggest the utility of consensus approaches as opposed to classical point estimates when we are interested in the topology of the phylogeny.

**Figure S1:**
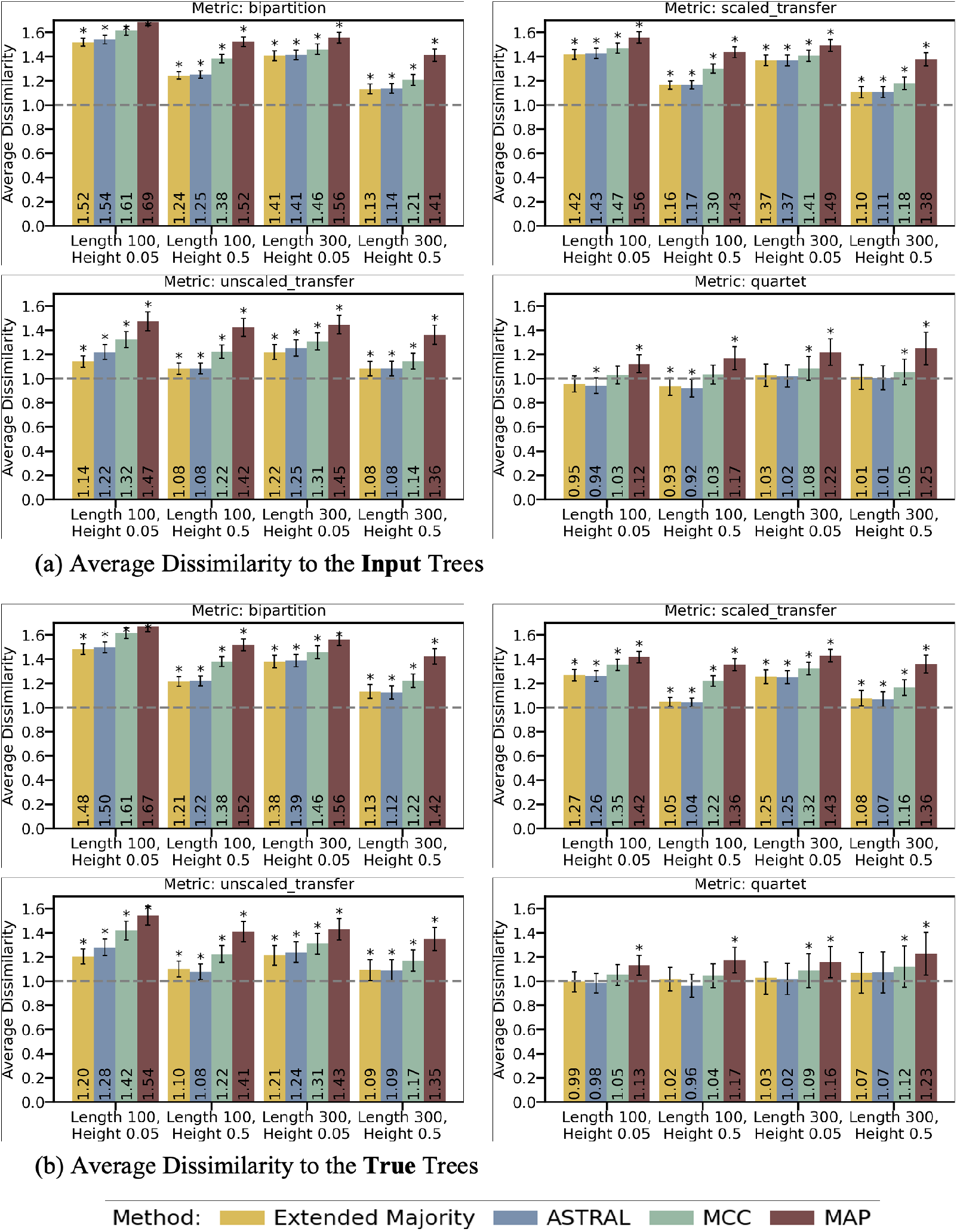
Performance of highly resolved trees in the Bayesian simulation experiment, in comparison to the majority rule (horizontal dashed line, Y = 1.0): (a) Average dissimilarities to the input trees for 100 datasets, (b) Average dissimilarities to the true trees for 100 datasets. The error bars are 95% confidence intervals, and asterisks indicate p-values (*p <* 0.01) based on paired t-tests of differences between each estimator and the majority-rule consensus tree. The extended-majority tree was fully resolved in all replicates except in the Length 100, Height 0.05 setting, where it was fully resolved in 70 of 100 replicates. The corresponding majority-rule baseline values are reported in Tables S1 and S2.

**Figure S2:**
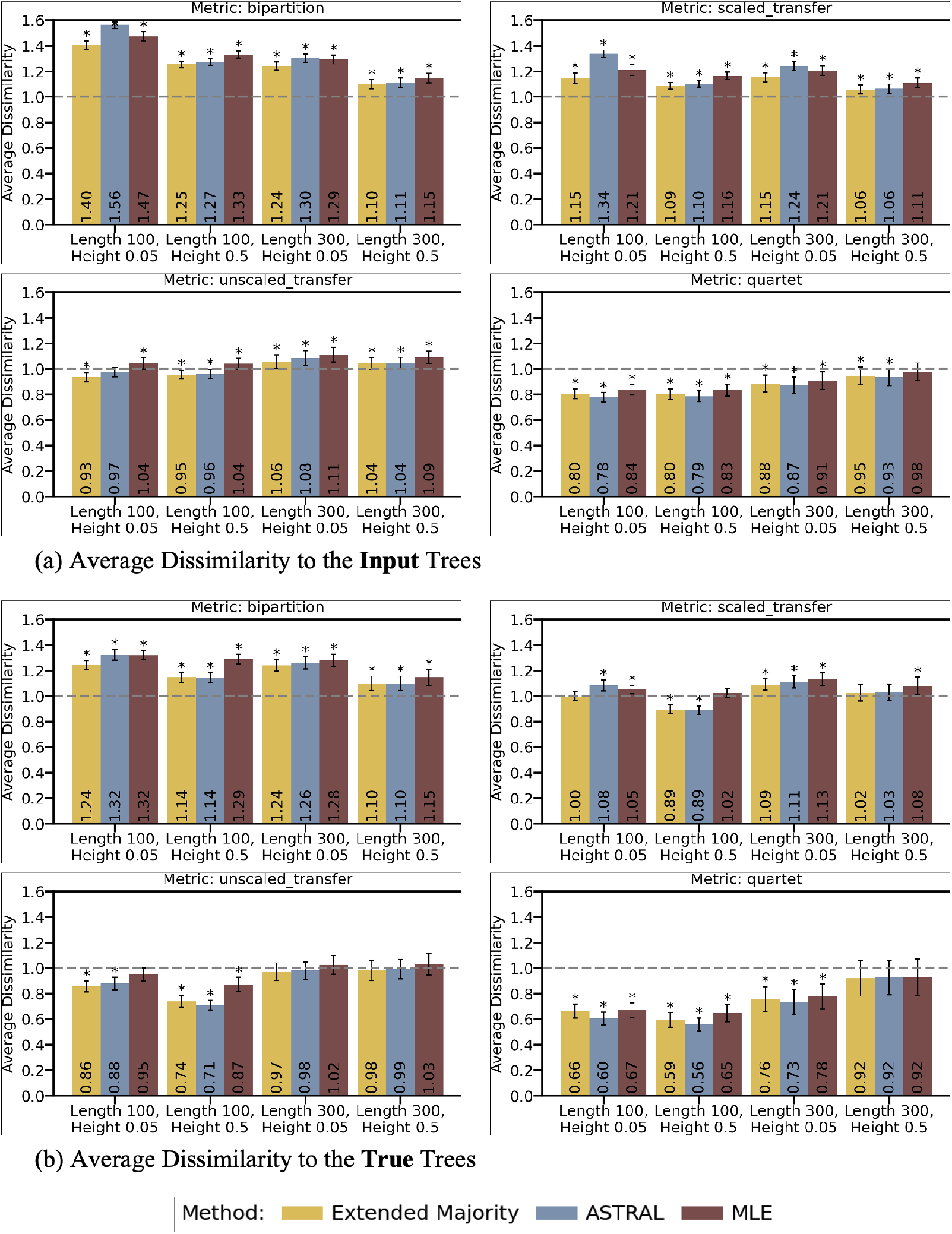
Performance of highly resolved trees in the bootstrap simulation experiment, in comparison to the majority rule (horizontal dashed line, Y = 1.0): (a) Average dissimilarities to the input trees for 100 datasets, (b) Average dissimilarities to the true trees for 100 datasets. The error bars are 95% confidence intervals, and asterisks indicate p-values (*p <* 0.01) based on paired t-tests of differences between each estimator and the majority-rule consensus tree. Unlike in the Bayesian experiment, extended majority was not always fully resolved: it was fully resolved in 2, 72, 17, and 96 of 100 replicates for Length 100/Height 0.05, Length 100/Height 0.5, Length 300/Height 0.05, and Length 300/Height 0.5, respectively. The corresponding majority-rule baseline values are reported in Tables S3 and S4.

### D.3 Average branch-support values in the simulation experiments

Tables S7–S9 report the average frequency-based and transfer-based support values. As expected, the majority-rule consensus generally has the highest average support because it contains fewer branches. The proposed consensus trees nevertheless retain substantially higher average support than the highly resolved alternatives. Among the proposed methods, the scaled-transfer consensus generally has slightly higher average support than the unscaled-transfer and quartet consensuses.

**Table S7:** Average support metrics in the Bayesian simulation (FS: frequency support, TS: transfer support).

| Method | Length 100 |  | Length 100 |  | Length 300 |  | Length 300 |  |
| --- | --- | --- | --- | --- | --- | --- | --- | --- |
|  | Height 0.05 |  | Height 0.5 |  | Height 0.05 |  | Height 0.5 |  |
|  | FS | TS | FS | TS | FS | TS | FS | TS |
| Majority | 0.86 | 0.93 | 0.87 | 0.92 | 0.92 | 0.96 | 0.93 | 0.96 |
| Scaled Transfer | 0.80 | 0.91 | 0.83 | 0.91 | 0.91 | 0.96 | 0.92 | 0.96 |
| Unscaled Transfer | 0.80 | 0.90 | 0.83 | 0.91 | 0.91 | 0.96 | 0.91 | 0.96 |
| Quartet | 0.79 | 0.90 | 0.80 | 0.89 | 0.90 | 0.95 | 0.91 | 0.95 |
| Extended Majority | 0.48 | 0.69 | 0.69 | 0.83 | 0.68 | 0.81 | 0.85 | 0.92 |
| ASTRAL | 0.47 | 0.69 | 0.69 | 0.83 | 0.68 | 0.82 | 0.85 | 0.92 |
| MCC | 0.45 | 0.70 | 0.65 | 0.82 | 0.67 | 0.81 | 0.84 | 0.92 |
| MAP | 0.42 | 0.68 | 0.62 | 0.80 | 0.65 | 0.80 | 0.81 | 0.90 |

**Table S8:** Support metrics in the bootstrap simulation (FBP: Felsenstein’s bootstrap proportion, TBE: transfer bootstrap expectation).

| Method | Length 100 |  | Length 100 |  | Length 300 |  | Length 300 |  |
| --- | --- | --- | --- | --- | --- | --- | --- | --- |
|  | Height 0.05 |  | Height 0.5 |  | Height 0.05 |  | Height 0.5 |  |
|  | FBP | TBE | FBP | TBE | FBP | TBE | FBP | TBE |
| Majority | 0.82 | 0.89 | 0.80 | 0.87 | 0.83 | 0.91 | 0.88 | 0.93 |
| Scaled Transfer | 0.73 | 0.85 | 0.74 | 0.85 | 0.81 | 0.90 | 0.86 | 0.93 |
| Unscaled Transfer | 0.71 | 0.85 | 0.72 | 0.84 | 0.80 | 0.90 | 0.86 | 0.92 |
| Quartet | 0.66 | 0.82 | 0.67 | 0.81 | 0.77 | 0.88 | 0.84 | 0.92 |
| Extended Majority | 0.52 | 0.75 | 0.59 | 0.78 | 0.67 | 0.83 | 0.81 | 0.90 |
| ASTRAL | 0.47 | 0.70 | 0.59 | 0.77 | 0.65 | 0.81 | 0.81 | 0.90 |
| MLE | 0.49 | 0.74 | 0.57 | 0.77 | 0.66 | 0.83 | 0.80 | 0.90 |

**Table S9:** Average support metrics in the Bayesian benchmark (FS: frequency support, TS: transfer support).

| Method | Coal320 |  | Yule400 |  |
| --- | --- | --- | --- | --- |
|  | FS | TS | FS | TS |
| Majority | 0.89 | 0.95 | 0.92 | 0.96 |
| Scaled Transfer | 0.85 | 0.95 | 0.91 | 0.96 |
| Unscaled Transfer | 0.85 | 0.94 | 0.91 | 0.95 |
| Quartet | 0.82 | 0.94 | 0.90 | 0.95 |
| Extended Majority | 0.70 | 0.88 | 0.83 | 0.91 |
| CCD0-MAP | 0.70 | 0.88 | 0.83 | 0.91 |
| ASTRAL | 0.70 | 0.88 | 0.83 | 0.91 |
| MCC | 0.65 | 0.85 | 0.80 | 0.90 |

### D.4 Branch-depth distributions

Figures S3 and S4 show the distributions of internal-branch depths. Across the simulation experiments and Bayesian benchmarks, the proposed consensus methods have broadly similar depth profiles to the input trees, with no systematic concentration of branches in a particular depth class.

**Figure S3:**
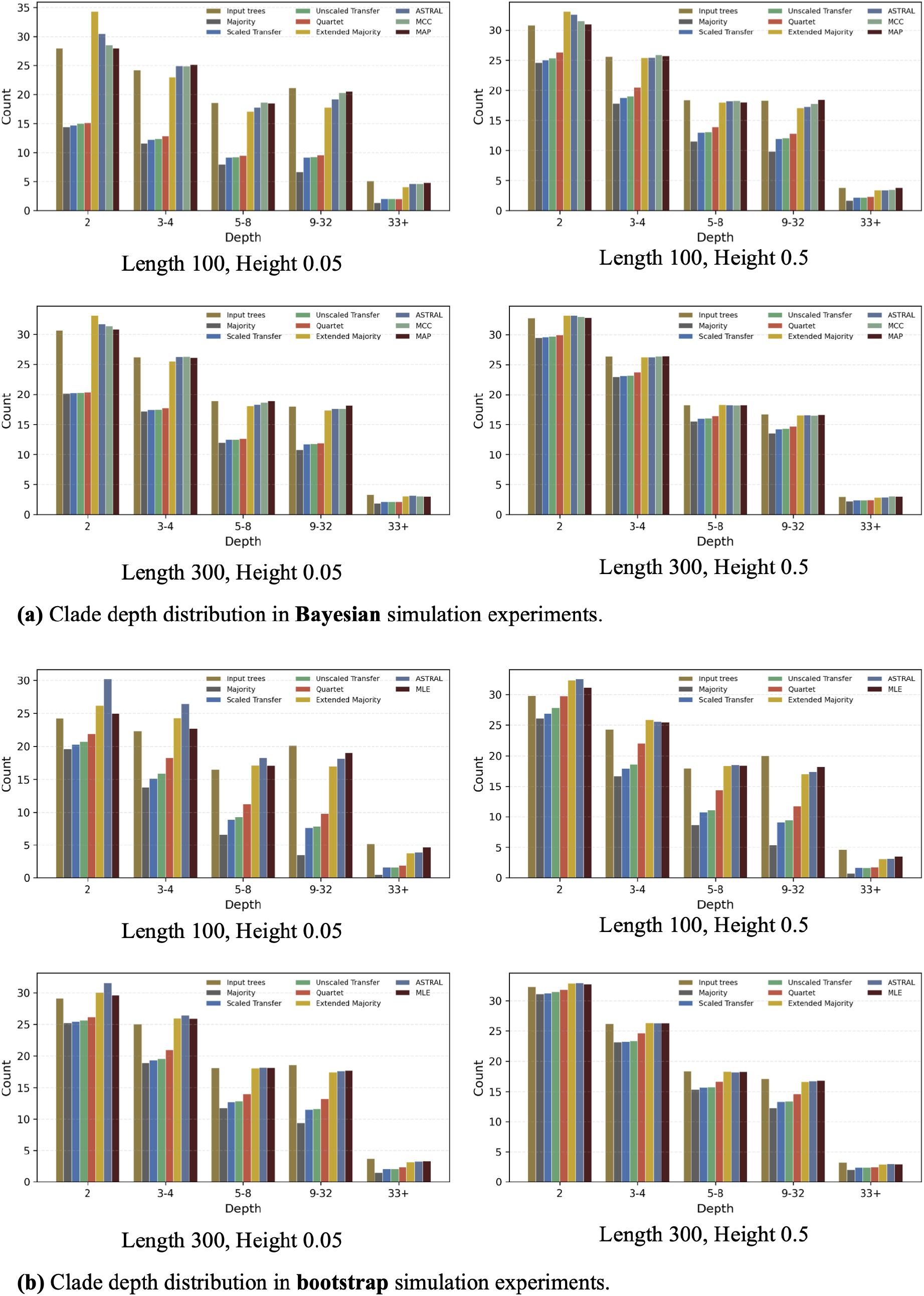
Internal-branch-depth distributions in the simulation experiments: (a) Bayesian analysis and (b) bootstrap analysis. Counts are averaged over the 100 replicate datasets.

**Figure S4:**
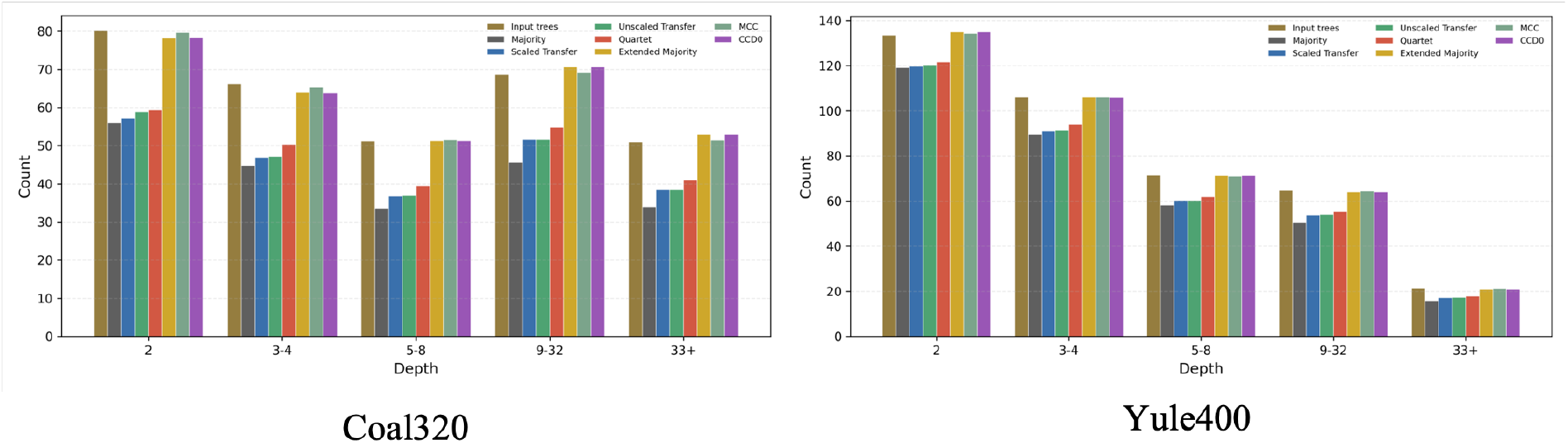
Internal-branch-depth distributions in the Bayesian benchmark experiments: Coal320 (left) and Yule400 (right). Counts for the consensus trees are averaged over the 100 replicate datasets. Again, we do not observe any strong bias for the proposed consensus methods.

### D.5 Bipartition overlap with the majority-rule consensus

Table S10 compares the internal bipartitions of each estimated tree with those of the majority-rule consensus. The proposed methods generally retain most majority-rule bipartitions while adding substantially more branches. In the simulation and Bayesian benchmark experiments, the mean number of majority-rule bipartitions absent from the transfer-based consensuses is at most 1.17, and the corresponding number for the quartet consensus is at most 0.11. In the EBOV analysis, only one to three of the 686 majority-rule bipartitions are absent from the proposed consensus trees. The same pattern is observed for the mammal and HIV datasets, although the overlap is somewhat lower for the full mammal dataset: the scaled- and unscaled-transfer consensuses miss 10 and 8 majority-rule bipartitions, respectively. However, this largely reflects the use of the MLE tree as the starting tree for pruning, since the MLE tree itself already lacks 7 of the majority-rule bipartitions. For HIV, the scaled- and unscaled-transfer consensuses retain 1321 and 1323 of the 1344 majority-rule bipartitions, respectively. Thus, overall, the increased resolution of the proposed methods is obtained predominantly by adding branches rather than by replacing majority-rule branches.

**Table S10:**
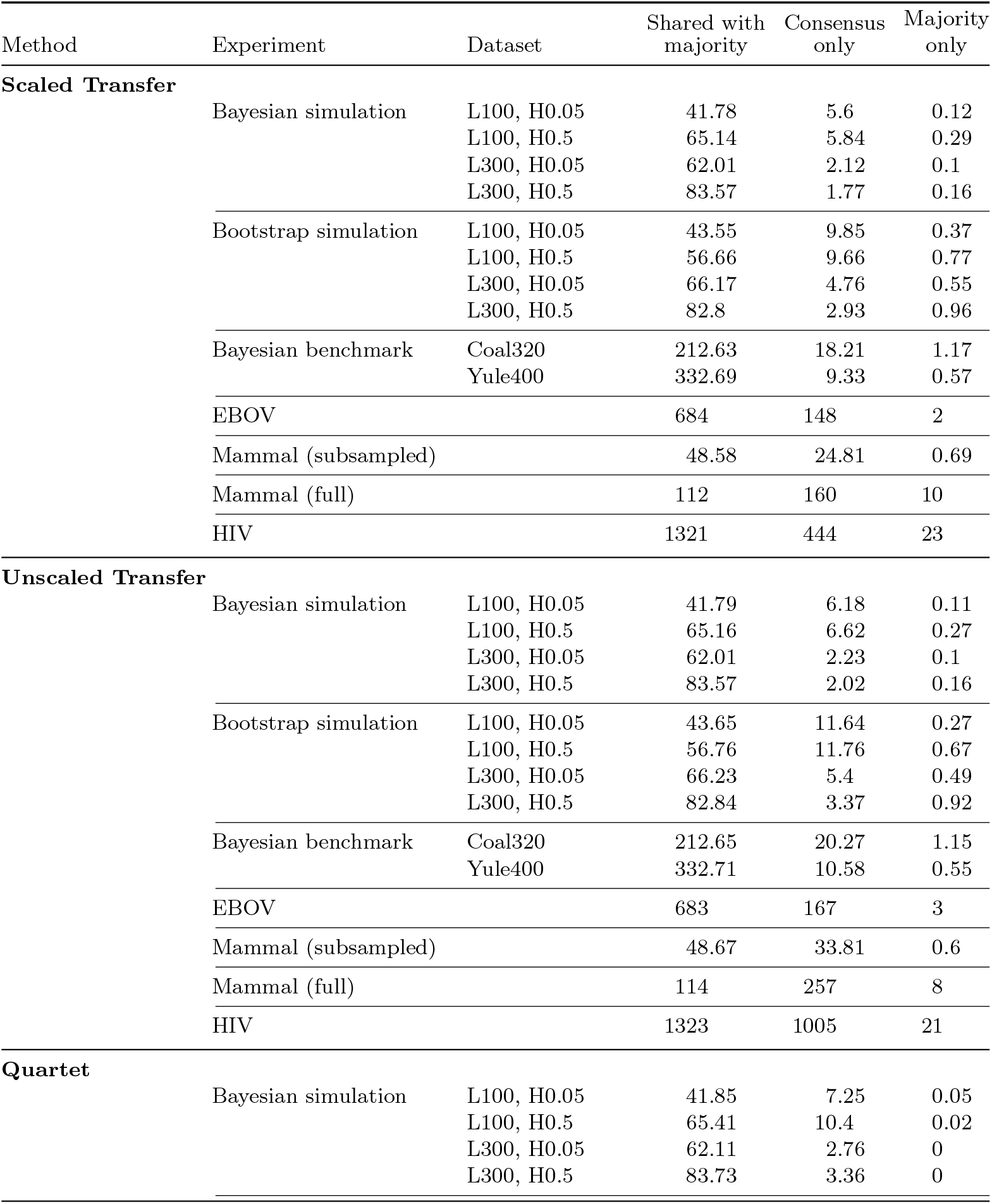
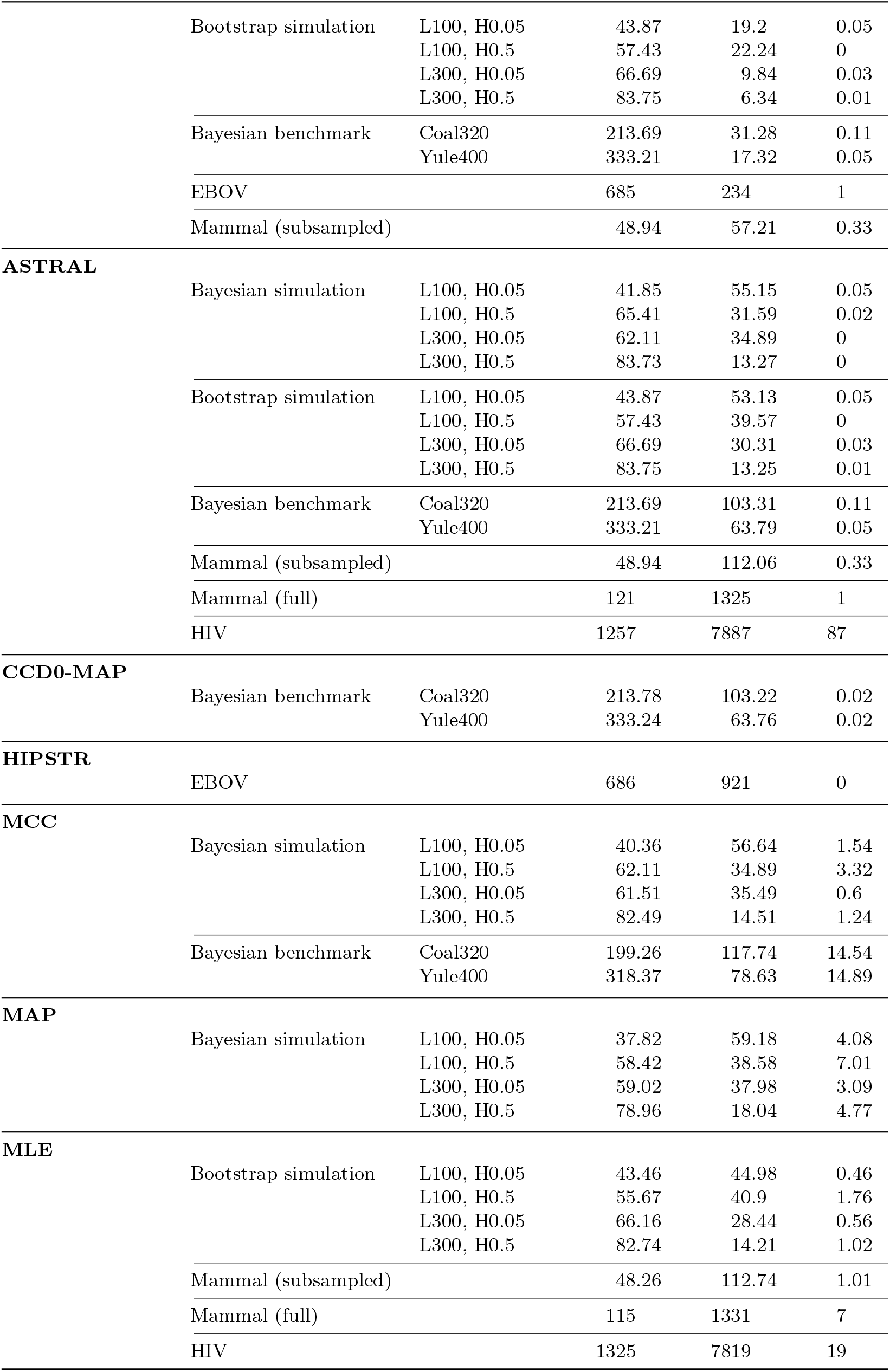
Comparison of the internal bipartitions in each estimated tree with those of the majority-rule consensus. “Shared with majority” denotes bipartitions present in both trees, “Consensus only” denotes bipartitions present only in the estimated tree, and “Majority only” denotes bipartitions present only in the majority-rule consensus. Values are averages over replicates when applicable.

## E User guide for the PhyloCRISP software

PhyloCRISP provides a graphical user interface (GUI) that enables users to generate consensus trees from a collection of phylogenetic trees in Newick format (Fig. S5). The interface supports the three proposed consensus methods (Scaled Transfer, Unscaled Transfer, and Quartet) as well as the classical Majority-Rule consensus. The workflow consists of loading input trees, selecting a consensus method and optimization strategy, optionally specifying a starting tree, generating the consensus tree, and exporting the result.

### Availability and download

Executable versions of PhyloCRISP for macOS, Linux, and Windows are available for download from the project GitHub repository https://github.com/yukiregista/PhyloCRISP. The same repository also provides the complete source code.

### Input trees

Users may paste Newick-formatted trees directly into the *Input Trees* panel or click *Browse Input Trees File* to load a Newick file. All trees must correspond to the same taxon set. For large files, the GUI may automatically hide the displayed content for performance reasons; this does not affect loading or computation.

### Consensus method

Under *Consensus Method*, users can choose one of the following:

- Scaled Transfer
- Unscaled Transfer
- Quartet
- Majority Rule

### Optimization strategy

For the three proposed methods, an optimization strategy must be specified:

- **Add and Prune**: The starting tree is optional. If not provided, the majority-rule consensus tree is used by default.
- **Prune Only**: A starting tree is required.

For the Quartet method, only the *Prune Only* strategy is currently available.

### Starting tree

If required, a starting tree can be entered directly in the *Starting Tree* panel or loaded via *Browse Starting Tree File*. The starting tree must be provided in Newick format and must use the same taxon set as the input trees.

### Generating and exporting the consensus tree

An output filename may be specified in the *Output Filename* field, and an output directory may also be selected. Clicking *Generate Consensus Tree* starts the computation. The resulting consensus tree is displayed in the *Consensus Tree* panel and can be exported using *Save Output*. The *Stop* button allows interruption of a running computation.

Three Newick trees are generated: one without support values, one with frequency support (FS, or FBP) values, and one with transfer support (TS, or TBE) values. These output trees are returned in standard Newick format and can be visualized with common phylogenetic tree viewers.

#### Practical guidance

- For the *Prune Only* strategy, we recommend using a fully resolved or nearly fully resolved starting tree (e.g., MLE, Extended Majority, or ASTRAL-IV).
- The *Add and Prune* strategy is more computationally demanding than *Prune Only* and may be impractical for datasets with several hundred taxa or more.
- The *Prune Only* strategy is computationally efficient for transfer-based dissimilarities. In practice, it can handle datasets with thousands of taxa (e.g., more than 9,000 leaves and 1,000 input trees) within moderate runtime (typically under one hour, depending on hardware).
- Computational time depends not only on tree size and number of input trees but also on their variability. Highly discordant input trees generally increase optimization time.

**Figure S5:**
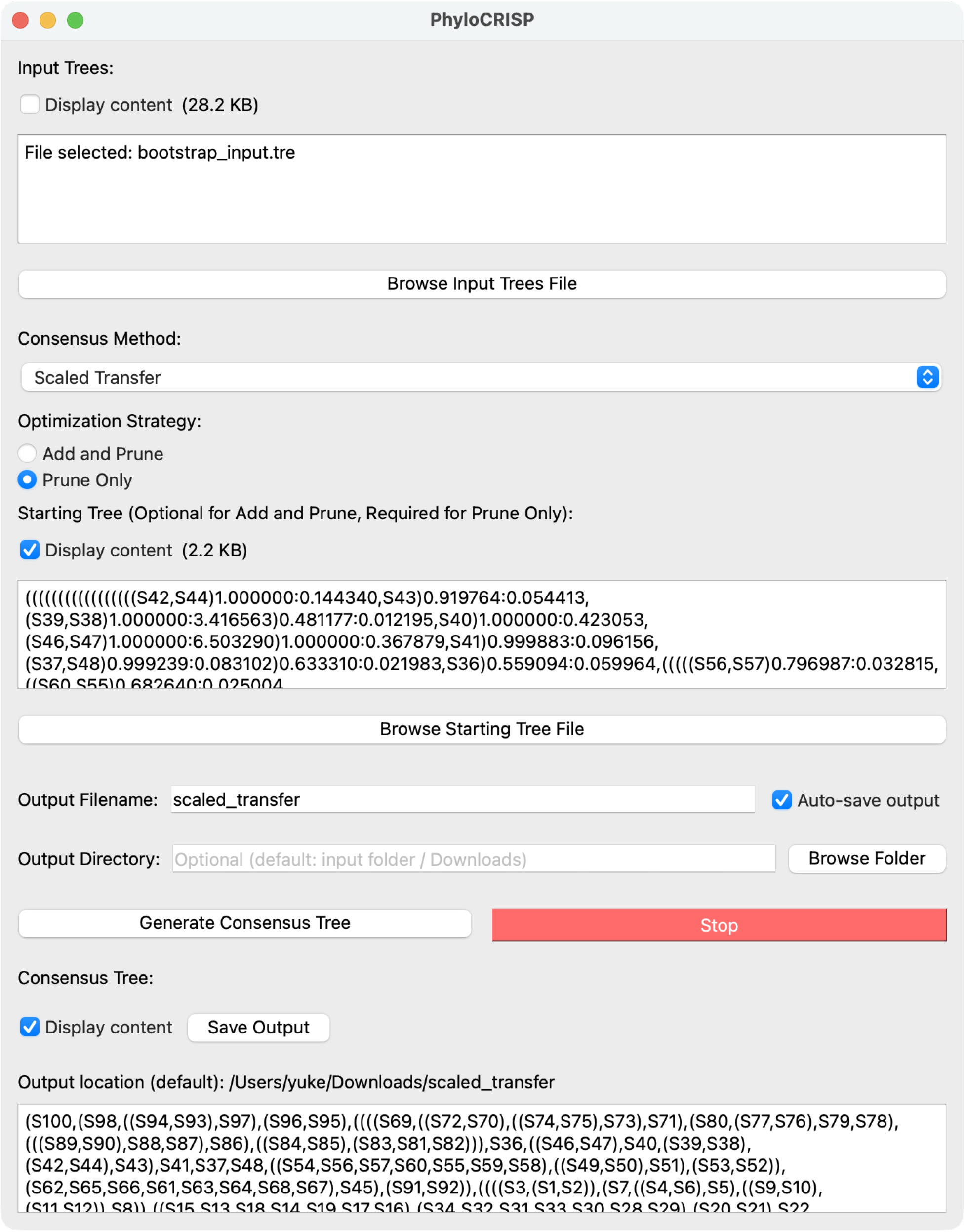
Screenshot of PhyloCRISP GUI.

## F Computational resource benchmark

To provide a practical indication of computational requirements, we measured wall-clock time and peak memory usage for the consensus methods on five shared replicates of the Yule400 dataset using our laptop (MacBook Pro, Apple M2 Pro, 32 GB memory). Since computational performance is not the primary focus of this study, these results are intended as representative resource requirements rather than a comprehensive software benchmark.

The transfer-based PhyloCRISP methods were computationally inexpensive. Using pruning only, the scaled- and unscaled-transfer methods required approximately 30 seconds per replicate, while the Add and Prune strategy required approximately 1.5 minutes. Quartet-based pruning was substantially more computationally demanding, requiring approximately 11 minutes. ASTRAL-IV, which is also based on quartets, required approximately 66 minutes on the same datasets using a single thread.

The empirical analyses show that both approaches can nevertheless be applied to large datasets. For the full EBOV dataset (1,610 taxa and 1,000 posterior trees), quartet-based pruning from the ASTRAL-IV tree required approximately 9 hours on our laptop. For the HIV dataset, comprising 1,000 bootstrap trees with 9,147 taxa, the transfer-based pruning analyses required approximately 20 minutes on the same laptop.

For comparison, generation of the input posterior or bootstrap tree collections is generally much more computationally demanding than the subsequent consensus calculation. The original BEAST2 analyses underlying the Yule400 benchmark required on the order of tens of hours per replicate with a single thread, while maximum-likelihood bootstrapping requires repeated tree inference.

For the HIV dataset, ASTRAL-IV itself was also computationally demanding. Using ASTER on the NIG supercomputer, it required approximately 214 hours (8.9 days) with 64 threads requested and reduced search settings (`-r 1 -s 1` specifying one initial search round and one subsampling and exploration round respectively; defaults: `-r 4 -s 4`).

**Table S11:** Computational resource usage on the Yule400 dataset. Values are means over five shared replicates. Time is wall-clock time in seconds, and memory is peak RSS in MB. In the command column, <trees> denotes the input collection of posterior trees and output-file options are omitted. Commands retain options relevant to the method or computational setting. PhyloCRISP 0.0.3 benchmarks used the installed public CLI.

| Method | Software | Command | Time (s) | Memory (MB) |
| --- | --- | --- | --- | --- |
| **PhyloCRISP consensus methods** |  |  |  |  |
| Scaled Transfer (prune) | PhyloCRISP 0.0.3 | `phylocrisp-cli --input-trees <trees> --method scaled_transfer --strategy prune --starting-tree astral.tre` | 25.37 | 911 |
| Unscaled Transfer (prune) | PhyloCRISP 0.0.3 | `phylocrisp-cli --input-trees <trees> --method unscaled_transfer --strategy prune --starting-tree astral.tre` | 25.59 | 916 |
| Scaled Transfer (add/prune) | PhyloCRISP 0.0.3 | `phylocrisp-cli --input-trees <trees> --method scaled_transfer --strategy add_and_prune --starting-tree mr.tre` | 99.53 | 1221 |
| Unscaled Transfer (add/prune) | PhyloCRISP 0.0.3 | `phylocrisp-cli --input-trees <trees> --method unscaled_transfer --strategy add_and_prune --starting-tree mr.tre` | 103.10 | 1221 |
| Quartet (prune) | PhyloCRISP 0.0.3 | `phylocrisp-cli --input-trees <trees> --method quartet --strategy prune --starting-tree astral.tre` | 649.34 | 277 |
| **Other consensus software** |  |  |  |  |
| Majority Rule | IQ-TREE 2.3.6 | `iqtree2 -con <trees> -minsup 0.5` | 1.40 | 285 |
| Extended Majority | IQ-TREE 2.3.6 | `iqtree2 -con <trees> -minsup 0` | 1.47 | 285 |
| ASTRAL-IV | ASTER 1.23 | `astral4 -i <trees> -t 1` | 3959.54 | 1100 |
| CCD0-MAP | BEAST 2.7.7 | `treeannotator -topology CCD0 -burnin 0 <trees> <out>` | 18.50 | 2797 |
| CCD1-MAP | BEAST 2.7.7 | `treeannotator -topology CCD1 -burnin 0 <trees> <out>` | 18.92 | 3169 |
| MCC | BEAST 2.7.7 | `treeannotator -topology MCC -burnin 0 <trees> <out>` | 19.37 | 3090 |
| HIPSTR | BEAST X 10.5.0 | `treeannotator -type hipstr -heights mean -burninTrees 0 -threads 1 <trees> <out>` | 3.12 | 630 |

